# Structure of an open K_ATP_ channel reveals tandem PIP_2_ binding sites mediating the Kir6.2 and SUR1 regulatory interface

**DOI:** 10.1101/2023.08.01.551546

**Authors:** Camden M. Driggers, Yi-Ying Kuo, Phillip Zhu, Assmaa ElSheikh, Show-Ling Shyng

## Abstract

ATP-sensitive potassium (K_ATP_) channels, composed of four pore-lining Kir6.2 subunits and four regulatory sulfonylurea receptor 1 (SUR1) subunits, control insulin secretion in pancreatic β-cells. K_ATP_ channel opening is stimulated by PIP_2_ and inhibited by ATP. Mutations that increase channel opening by PIP_2_ reduce ATP inhibition and cause neonatal diabetes. Although considerable evidence has implicated a role for PIP_2_ in K_ATP_ channel function, previously solved open-channel structures have lacked bound PIP_2_, and mechanisms by which PIP_2_ regulates K_ATP_ channels remain unresolved. Here, we report cryoEM structure of a K_ATP_ channel harboring the neonatal diabetes mutation Kir6.2-Q52R, bound to natural C18:0/C20:4 long-chain PI(4,5)P_2_ in open conformation. The structure reveals two adjacent PIP_2_ molecules between SUR1 and Kir6.2. The first PIP_2_ binding site is conserved among PIP_2_-gated Kir channels. The second site forms uniquely in K_ATP_ at the interface of Kir6.2 and SUR1. Functional studies demonstrate both binding sites determine channel activity. Kir6.2 pore opening is associated with a twist of the Kir6.2 cytoplasmic domain and a rotation of the N-terminal transmembrane domain of SUR1, which widens the inhibitory ATP binding pocket to disfavor ATP binding. The open conformation is particularly stabilized by the Kir6.2-Q52R residue through cation-π bonding with SUR1 - W51. Together, these results uncover the cooperation between SUR1 and Kir6.2 in PIP_2_ binding and gating, explain the antagonistic regulation of K_ATP_ channels by PIP_2_ and ATP, and provide the mechanism by which Kir6.2-Q52R stabilizes an open channel to cause neonatal diabetes.

## Introduction

The K_ATP_ channel expressed in pancreatic islet β-cells is a principal homeostatic regulator of insulin secretion^1–3^. Composed of four pore-forming Kir6.2 and four regulatory sulfonylurea receptor 1 (SUR1) subunits^4,5^, K_ATP_ channels influence the resting membrane potential, thus action potential firing, Ca^2+^ entry, and insulin secretion. The activity of K_ATP_ channels is dynamically regulated by glucose changes via intracellular ATP and MgADP, which bind to inhibitory and stimulatory sites to close and open the channel, respectively^6,7^. This enables insulin secretion to follow fluctuations in glucose concentrations. The central role of K_ATP_ channels in insulin secretion and glucose homeostasis is underscored by dysregulated insulin secretion and blood glucose in humans bearing K_ATP_ mutations: loss-of-function mutations cause congenital hyperinsulinism characterized by persistent insulin secretion despite life-threatening hypoglycemia^8,9^; conversely, gain-of-function channel mutations result in neonatal diabetes due to insufficient insulin secretion^10^.

In addition to adenine nucleotides-dependent regulation, membrane phosphoinositides participate in the gating activation of K_ATP_ channels^11,12^, as they do all other eukaryotic Kir channels^13^. PI(4,5)P_2_ (denoted as PIP_2_ unless specified) is the most abundant phosphoinositide in the plasma membrane^14^, with PIP_2_ containing arachidonoyl (C20:4) and stearoyl (C18:0) acyl chains being the most dominant species in mammalian cells^15^. In inside-out patch-clamp recordings, application of C18:0/C20:4 long-chain PIP_2_ to the cytoplasmic side of the membrane patch resulted in stable increases in K_ATP_ channel open probability (*P_o_*) and a gradual decrease in channel sensitivity to ATP inhibition; whereas scavenging membrane PIP_2_ with poly-lysine decreases channel *P_o_*^11,12^. Because increasing channel *P_o_* by PIP_2_ also decreases channel sensitivity to ATP inhibition^11,12^, a joint regulatory mechanism involving allosteric or binding competition between the two ligands has seemed likely^16^. In particular, the effect of PIP_2_ on K_ATP_ channels has been assumed to derive from a putative conserved binding site in Kir6.2 having a position homologous to the PIP_2_ binding site previously identified in crystal and cryoEM structures of Kir2 and Kir3^17–20^. Uniquely however, gating of Kir6.2 by PIP_2_ is strongly modulated by its regulatory subunit SUR1. Compared to the fully assembled K_ATP_ channels, Kir6.2 channels lacking SUR1 partners display more than 10-fold lower *P_o_*^11,16,21^, which reflects reduced PIP_2_ binding or gating, and are less sensitive to PIP_2_ stimulation^11,16^. Of particular interest, the SUR1 N-terminal transmembrane domain, TMD0, which interfaces with Kir6.2^22-24^, is sufficient to confer the high *P_o_* of wild-type (WT) channels^25,26^, implicating a crucial role of SUR1-TMD0 in controlling Kir6.2 response to PIP_2_.

To date, cryoEM structures of K_ATP_ channels determined for a variety of liganded and mutational states have yielded a structural spectrum encompassing open, closed, and inactivated conformations (reviewed in^6,7,27^). Careful comparisons have suggested mechanisms by which channel activity is regulated by the SUR subunit and nucleotides. However, confusion has grown regarding PIP_2_ binding and gating. Two recently reported open- and pre-open mutant K_ATP_ channel cryoEM structures lack discernible PIP_2_ at the putative binding site^28,29^, even when excess synthetic short-chain diC8-PI(4,5)P_2_ was added to the sample in the case of the pre-open structure^30^. Furthermore, purified WT K_ATP_ channels reconstituted in lipid bilayer s lacking PIP_2_ showed single channel openings^28^. These reports suggest that PIP_2_ binding may be inessential for open channel transition^28^. Thus, notwithstanding robust data indicating PIP_2_ involvement in physiological channel activation^11,31,32^, how PIP_2_ binds and modulates K_ATP_ channels remains enigmatic.

Our approach to understanding K_ATP_ channel activation and PIP_2_ binding involved determining channel structure bound to PIP_2_ by (1) incubating membranes expressing K_ATP_ channels with long-chain PIP_2_ known to stably activate the channel^11,12,33^, and (2) employing a Kir6.2 variant containing the neonatal diabetes-causing mutation Q52R (denoted Kir6.2^Q52R^)^34,35^. Kir6.2^Q52R^ markedly increases channel *P_o_* and causes a 20-fold decrease in channel sensitivity to ATP inhibition^34,35^. Higher *P_o_* could result from enhanced PIP_2_ binding or its effect on gating. Intriguingly, the effect of this mutation on channel gating is SUR1-dependent. When Kir 6.2 was manipulated to express at the cell surface without co-assembly with SUR1 (by deleting the Arg-Lys-Arg endoplasmic reticulum retention signal at the C-terminal region of Kir6.2), introducing the Q52R mutation had no effect on the *P_o_* and ATP sensitivity^36^. The SUR1-dependent over-activity, imparted by the Kir6.2-Q52R mutation, indicated the possibility to probe the regulatory interaction between SUR1 and membrane PIP_2_ on Kir6.2 gating.

Here, we present the cryoEM structure of the K_ATP_ channel harboring the neonatal diabetes Q52R mutation (referred to as the SUR1/Kir6.2^Q52R^ channel hereinafter) bound to C18:0/C20:4 long-chain PI(4,5)P_2_ in an open conformation. Surprisingly, two PIP_2_ molecules bind in tandem in the open channel at the interface between Kir6.2 and SUR1. A first PIP_2_ molecule bound within the site that is conserved among Kir channels. A second PIP_2_ molecule was at a novel adjacent position, stacked within the regulatory interface between Kir6.2 and SUR1 TMDs. The open channel conformation is further stabilized by direct interaction between Q52R of Kir6.2 and W51 located in TMD0 of SUR1. Accounting for decades of evidence from functional perturbation studies, the PIP_2_-bound SUR1/Kir6.2^Q52R^ K_ATP_ structure unveils the agonist binding of PIP_2_ in K_ATP_ channels, revealing that in addition to the ancestral Kir PIP_2_ site, the incorporation of the regulatory SUR1 subunit creates a second regulatory PIP_2_ binding site. The results implicate a novel mechanism through which SUR1, by cooperating with Kir6.2 for PIP_2_ binding, stabilizes the K_ATP_ channel in an open state to control activity, and illuminate the molecular mechanism by which a human activating mutation in the K_ATP_ channel causes neonatal diabetes.

## Results

### Structure of an open SUR1/Kir6.2^Q52R^ K_ATP_ channel bound to PIP_2_

To purify the neonatal diabetes variant SUR1/Kir6.2^Q52R^ K_ATP_ channel, we co-expressed Kir6.2^Q52R^ and SUR1 as independent polypeptides by transducing adherent mammalian COSm6 cells with recombinant adenovirus packaged with genes for rat Kir6.2 ^Q52R^ and FLAG-tagged hamster SUR1 (see Methods). Prior functional experiments have shown that the activating effects of long-chain PIP_2_ are long-lasting and resistant to subsequent washout, indicating stable partition into the membrane^11,12,33^. By contrast, the effects of synthetic short - chain PIP_2_ such as diC8-PIP_2_ are highly reversible^33,37^. Accordingly, we prepared a membrane fraction of COSm6 cells expressing the mutant channel, and enriched the native membranes with brain PIP_2_, primarily containing C18:0/C20:4 long-chain PI(4,5)P_2_ (Avanti Polar Lipids), before detergent solubilization. We anticipated that addition of long-chain PIP_2_ prior to detergent extraction would stably incorporate PIP_2_ resistant to washout in subsequent purification steps to facilitate the capture of PIP_2_-bound channel structure. K_ATP_ channels were purified via the FLAG-tag on SUR1 and spotted on graphene oxide (GO)-coated grids for cryoEM (see Methods). To maximize channels in PIP_2_-bound open conformations, no ATP or ADP were added to the sample.

CryoEM micrographs showed mostly full SUR1/Kir6.2^Q52R^ K_ATP_ channel particles (Fig. S1a). 2D classes indicate an ordered Kir6.2 and SUR1-TMD0, with a relatively more blurred SUR1-ABC core especially the nucleotide binding domains (NBDs) (Fig. S1a). A non-uniform refinement with C4 symmetry using all good particles from 2D classification (70,638) in cryoSPARC resulted in an un-masked reconstruction of the full channel at 3.9 Å using the gold-standard Fourier shell correlation (GSFSC) cutoff of 0.143 (see Methods). The electron potential map shows relatively weak signal for TMD2 and NBD2 of SUR1 (Fig. S1), indicating higher mobility of these domains. Multiple rounds of *ab initio* reconstruction without symmetry imposed (Fig. S1a) yielded a best 3D class containing 14,115 particles, which following a C4 non-uniform refinement in cryoSPARC with auto-masking that excluded SUR1-NBD2 resulted in a final map reconstruction of 2.9 Å by GSFSC within the autoFSC mask (Figs. 1a, S1), or 3.3 Å resolution by GSFSC within a mask that includes the full K_ATP_ channel. Since the full channel map derived from the C4 non-uniform refinement has the best overall resolution, showing a highly ordered Kir6.2 tetramer core and SUR1-TMD0 with local resolutions < 2.9 Å, the map was used for model building (Figs. 1, S1, Table S1). The structure revealed bound PIP_2_ at the regulatory interface of Kir6.2 and SUR1 and an open Kir6.2 pore and molecular rearrangements at the SUR1-Kir6.2 interface critical to gating, as we describe in detail in the following sections.

**Figure 1.**
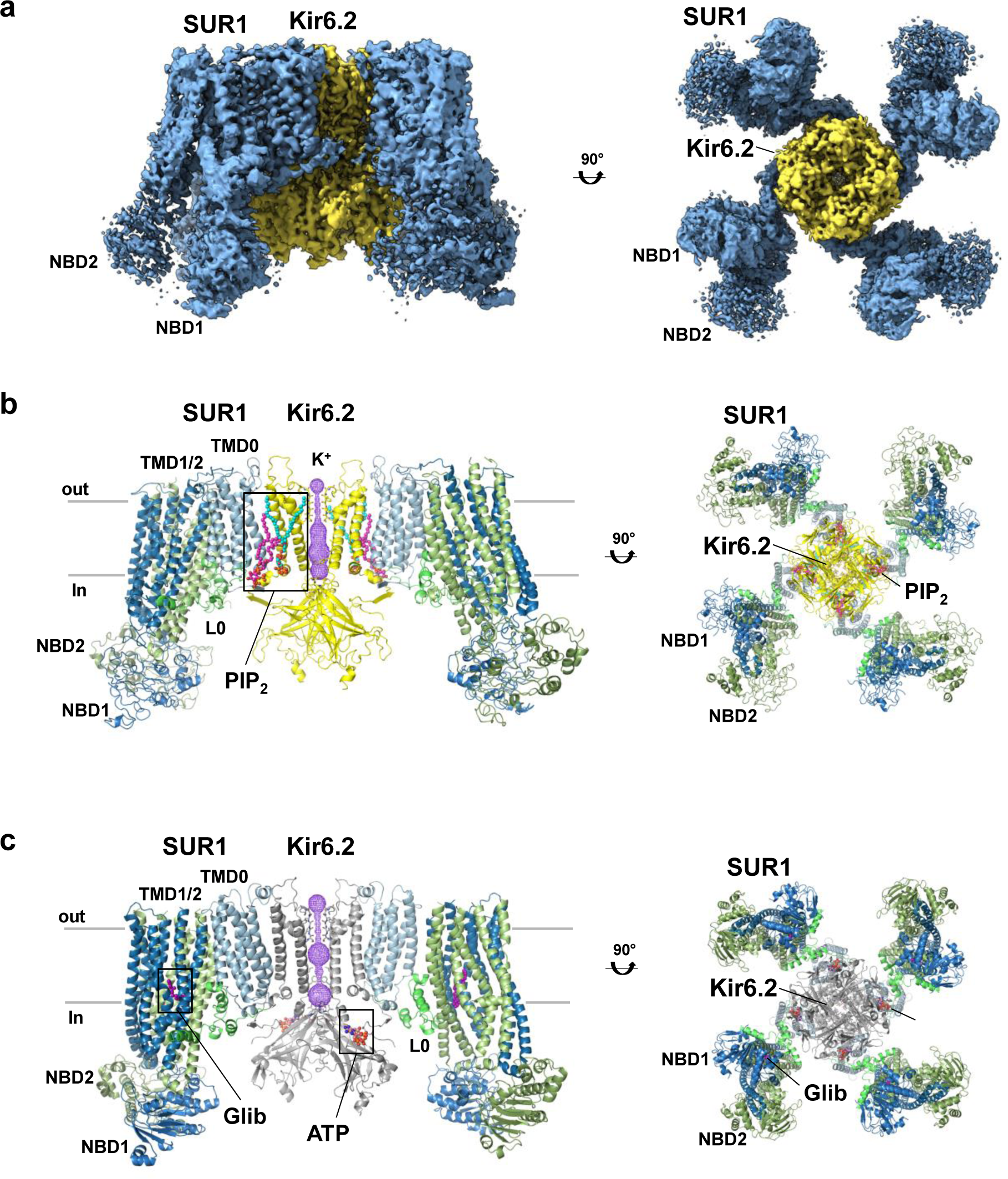
CryoEM structure of PIP_2_-bound, open SUR1/Kir6.2^Q52R^ K_ATP_ channel. **(a)** The 2.9 Å cryoEM map (the micelle not shown), from C4 non-uniform refinement of 14,115 particles, shown from the side and bottom (cytoplasmic view). **(b)** Structural model of the SUR1/Kir6.2^Q52R^ K_ATP_ channel in complex with PIP_2_. The potassium ion selectivity filter is shown as sticks with oxygens colored red, the K^+^ conduction pathway is shown as purple mesh. Only two Kir6.2 and two SUR1 subunits are shown in the side view for clarity. A view rotated 90° (cytoplasmic view) displays all four Kir6.2 subunits and all four SUR1 subunits. **(c)** Structural model of a closed WT SUR1/Kir6.2 K_ATP_ channel in complex with ATP and gliblenclamide (Glib) (PDB ID 6BAA), shown for comparison and rendered in PyMOL similar to (b), except for the Kir6.2 subunits which are colored grey.

Additional analysis of all particles from the 2D classes employed a strategy of symmetry expansion and focused 3D classification (using a mask of Kir6.2^Q52R^ tetramer plus one SUR1 subunit to circumvent the flexibility of the SUR1 subunits), revealing a spectrum of conformational classes with distinct Kir6.2-cytoplasmic domain (CTD) and SUR1 positions (Fig. S2). The two most extreme conformations include a dominant conformation at ∼3.3 Å ( no C4 symmetry imposed) that is nearly identical to the map derived from the C4 non-uniform refinement described above, and a minor conformation at 6.9 Å (Fig. S2a). Superimposition of the two extreme conformations revealed differences in the K^+^ pore path, rotation of the Kir6.2^Q52R^-CTD and distance of Kir6.2^Q52R^-CTD to the membrane (Fig. S2b-d), as well as the absence or presence of the cryoEM density corresponding to the N-terminal domain of Kir6.2^Q52R^ (the distal 30 amino acids, referred to as KNtp; Fig. S2e). Relative to the high resolution dominant class, the low resolution minor class shows a more constricted Kir6.2^Q52R^ pore, Kir6.2^Q52R^-CD clockwise rotated (viewed from the cytoplasmic side) and moved away from the membrane, and clear KNtp density in the central cleft of the SUR1-ABC core. Moreover, the SUR1-ABC core is rotated towards the Kir6.2^Q52R^ tetramer and moves downward from the outer membrane (Fig. S2c,d). These features highly resemble our previously published closed apo (no PIP_2_, nucleotides, or pharmacological inhibitors added) WT K_ATP_ channel structure (PDB ID 7UQR) ^38^, and indicate the minor conformation isolated from our sample represents an apo closed SUR1/Kir6.2^Q52R^ structure (and will be referred to as such hereinafter) even with the exogenously added long-chain PIP_2_.

### Tandem PIP_2_ binding at the interface of Kir6.2 and SUR1

The cryoEM map of PIP_2_-bound SUR1/Kir6.2^Q52R^ K_ATP_ channels showed lipid density corresponding to two adjacent PIP_2_ molecules occupying sites in a large pocket at the Kir6.2-SUR1 interface near the inner leaflet of the membrane bilayer ( Fig. 2a). The first PIP_2_ occupies the predicted PIP_2_ binding site of Kir6.2 based on comparisons with the homologous structures of Kir channels Kir2 and Kir3/GIRK bound to PIP_2_^17–20^ (colored cyan in all relevant figures). The second PIP_2_ binding site, which is occupied by a PIP_2_ with stronger map features (colored magenta in relevant figures) than the first PIP_2_, is nestled between SUR1 and the first PIP_2_ binding site and in contact with both Kir6.2 and SUR1 subunits (Fig. 2). Both PIP_2_ molecules were fit into the map by modelling as C18:0/C20:4 PI(4,5)P_2_ (Figs. 1b, 2b,c, S3a, b), which is the dominant form present in brain PIP_2_ (∼75%)^15^, and both PIP_2_ sites have map features for the phospho-head groups as well as long acyl chains that can be fit by a C18:0/C20:4 PI(4,5)P_2_ structure (monomer ID PT5) (Figs. 2b, c, S3b). Three other well-resolved lipid map features at the inner leaflet of the membrane bilayer were bound between SUR1-TMD0 and SUR1 - TMD1. These were best modeled by the common native phospholipids phosphatidylethanolamine and phosphatidylserine (Fig. S3c,d). The comparative fit of the different lipid species demonstrates that they can be distinguished using the reconstructed cryoEM maps (Fig. S3). Of note, lipid or detergent densities were also observed in the outer leaflet space (Fig. S3a,c); however, they were not sufficiently resolved to allow modeling.

**Figure 2.**
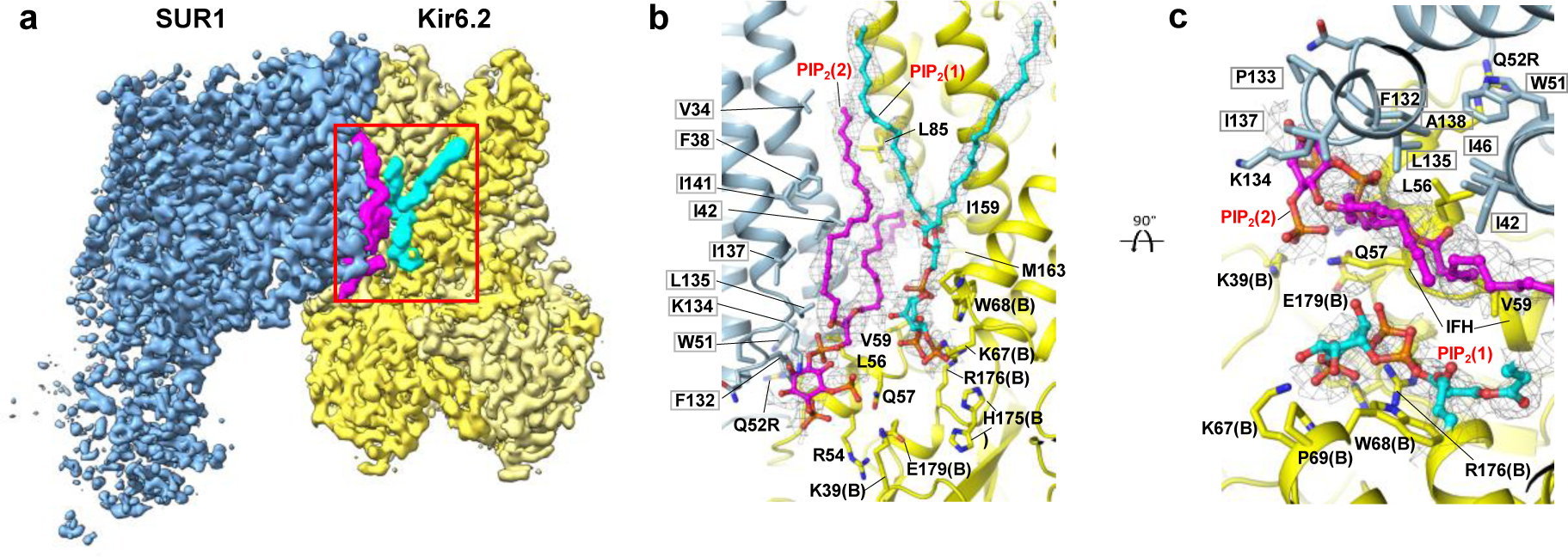
Two PIP_2_ binding sites at the interface of Kir6.2 and SUR1. **(a)** CryoEM map features of two PIP_2_ molecules colored in cyan and magenta (0.08V, 4.0σ contour), respectively. **(b)** Structural model of the PIP_2_ binding pocket (red boxed region in (a)) viewed from the side, with cryoEM density of PIP_2_ shown as grey mesh. Residues from both Kir6.2 (adjacent subunit denoted “B”) and SUR1 (grey outline) surrounding bound PIP_2_ molecules are labeled. **(c)** Structural model of the PIP_2_ binding pocket view from the top (extracellular side).

#### Structural analysis surrounding the PIP_2_ head groups

The head group of PIP_2_ at the first binding site forms polar interactions with surrounding residues (Fig. 2b, c). The phosphate groups are coordinated directly by Kir6.2 residues R176 and K67 while Kir6.2-W68 forms close interaction with the inositol ring, although K170, which is about 4.5 Å away and H175, which is about 5.5 Å away, are also in close proximity. Most of the head group-interacting residues are conserved in Kir2 and Kir3 channels with the exception of Kir6.2-P69 and H175, which in Kir2 and Kir3 are an arginine and a lysine, respectively and shown to coordinate PIP_2_ binding in Kir2 and Kir3^17–20^. The amino acid difference at these two positions would weaken Kir6.2-PIP_2_ interactions, consistent with the lower phosphoinositide head-group specificity compared to other Kir channels^33^. Interestingly, Kir6.2-H175 sidechain exhibited two rotamer positions in the PIP_2_-bound SUR1/Kir6.2^Q52R^ channel cryoEM density map: one oriented towards PIP_2_ in the conserved binding site and the other towards E179 in the same Kir6.2 subunit (Fig. 2b). Kir6.2-H175 has been previously implicated in acid-induced activation of K_ATP_ channels between pH 7.4 and 6.8 with a pK of 7.16^39^. Mutating Kir6.2-H175 to lysine, which mimics protonated state of H175, increased basal channel activity and abolished channel activation by pH, suggesting protonation of H175 favors channel opening ^39^. Moreover, mutating Kir6.2-E179 to Q greatly attenuated pH-induced channel activation, suggesting Kir6.2 - E179 also has a role in acid activation of K_ATP_ channels. Our structural observation that H175 sidechain has interaction with PIP_2_ head group in the conserved site and with Kir6.2-E179 is consistent with the aforementioned mutation-function correlation studies. Since our protein sample was prepared at pH∼7.5 (see Methods), which predicts H175 to be largely unprotonated based on the pK of acid activation of the channel of 7.16, it remains to be determined how protonation of H175 at lower pH that activates the channel alters interaction with PIP_2_ and E179 to stabilize channel opening.

The second PIP_2_ molecule sits in a novel site coordinated by both Kir6.2 and SUR1 subunits (Figs. 1b, 2). Here, the PIP_2_ head group interacts with surrounding residues including K39 and Q57 of Kir6.2, and N131, F132, P133, and K134 of SUR1. Previously we have noted that in closed K_ATP_ structures, SUR1-K134, located in intracellular loop (ICL)2 of SUR1 - TMD0, has its side chain directed towards cryoEM densities in the predicted PIP_2_ binding pocket tentatively assigned as phosphatidylserines^38^. Based on the observation, we mutated SUR1-K134 to alanine and found it reduced channel *P_o_*, which was recovered by adding exogenous PIP_2_^38^. Our PIP_2_-bound open structure now shows that SUR1-K134 is involved in PIP_2_ binding but at the novel second site, directly explaining the mutational study result.

#### Structural interactions with the acyl chains of PIP_2._

Most phospholipids present a range of molecular species comprising acyl chains of diverse length and saturation. However, mammalian PIPs show the predominance of a single hydrophobic backbone composed of arachidonoyl ( C20:4) and stearoyl (C18:0) chains^15^. In the open K_ATP_ channel structure determined with an enrichment of natural brain PIP_2_ in the membrane, we observed cryoEM densities corresponding to branched long acyl chains for both PIP_2_ molecules and modeled them to best fit the densities (Figs. 2b, S3a, S3b). The first PIP_2_ bound at the Kir-conserved site, mediates interactions between Kir6.2 subunits. While one chain is primarily associated with the outer TM helix of the Kir6.2 subunit that coordinates head group binding (modeled as the arachidonoyl chain), the other chain sits between the TM helicies of the adjacent Kir6.2 subunit (modeled as the stearoyl chain; Fig. 2b). CryoEM density at this conserved PIP_2_ binding pocket was previously reported by our group and others, but in those studies the map features lack lipid acyl tails and could not be ascertained as PIP_2_^30,38,40,41^ (Fig. S4a,b). The lipid tails of the second PIP_2_ molecule are sandwiched between SUR1 - TMD0 and Kir6.2-TMs, with one chain running along TM1 and TM4 of SUR1-TMD0 (modeled as the arachidonoyl chain) and the other between TM1 of SUR1-TMD0 and TMs of Kir6.2 (modeled as the stearoyl chain), and have non-polar interactions with SUR1 (V34, F38, I42, P45, I46, L135, I137, A138, I141) and Kir6.2 (L56, V59, I159) respectively (Fig. 2b, c). Long-chain PIP_2_ stably activates K_ATP_ channels^11,12^; in contrast, activation of channels by synthetic short-chain PIP_2_ such as diC8-PIP_2_ is readily reversible^33^. The extensive hydrophobic interactions between channel subunits and the long acyl chains of C18:0/C20:4 PI(4,5)P_2_ observed in our structure may contribute to the apparent increase in stability of channel activation by long-chain PIP_2_.

#### Functional role of the two PIP_2_ binding sites

To probe the functional role of the two PIP_2_ binding sites, we compared the PIP_2_ response of WT channels to channels containing the following mutations: Kir6.2^R176A^, which is predicted to weaken the first PIP_2_ binding site; SUR1^K134A^, which is expected to weaken the second PIP_2_ binding site; Kir6.2^R176A^ and SUR1^K134A^, which weaken both PIP_2_ binding sites (Fig. 3a). Exposure to brain-derived C18:0/C20:4 PI(4,5)P_2_ increased activity that is resistant to washout and decreased ATP inhibition over time in WT channels, as has been shown previously^11,12^. Similar effects of PIP_2_ were observed in all three mutant channels (Fig. 3a). However, the initial currents upon membrane excision, the extent of current increase upon PIP_2_ stimulation, and the PIP_2_ exposure time required for currents to reach maximum showed striking differences among the different channels (Fig. 3b-d). WT channels exhibited robust activity upon membrane excision into ATP-free solution, and currents increased by 1.48 ± 0.23-fold (Fig. 3a) that plateaued within one minute of PIP_2_ exposure. Perturbation of the second PIP_2_ binding site by SUR1^K134A^ resulted in channels that showed reduced initial currents (596.91 ± 218.23 pA vs. 1488.53 ± 348.86 pA of WT channels), which increased by 3.96 ± 0.47-fold in response to PIP_2_. Perturbation of the first PIP_2_ binding site by Kir6.2^R176A^ markedly reduced initial currents (42.44 ± 11.12 pA), which increased by 48.74 ± 18.25-fold after PIP_2_ exposure. Combining SUR1^K134A^ and Kir6.2^R176A^ yielded channels that showed barely detectable currents at patch excision (12.62 ± 4.34 pA) and a dramatic 299.65 ± 121.72-fold current increase by PIP_2_. All three mutants also required longer PIP_2_ exposure to reach maximum currents in the order of double mutant (419.33 ± 57.86 sec) > Kir6.2^R176A^ (298.57 ± 73.76 sec) > SUR1^K134A^ (193.83 ± 33.66 sec), compared to WT channels (33.17 ± 3.18 sec) (Fig. 3d). These results demonstrate that both PIP_2_ binding sites have functional roles in K_ATP_ channel activity.

**Figure 3.**
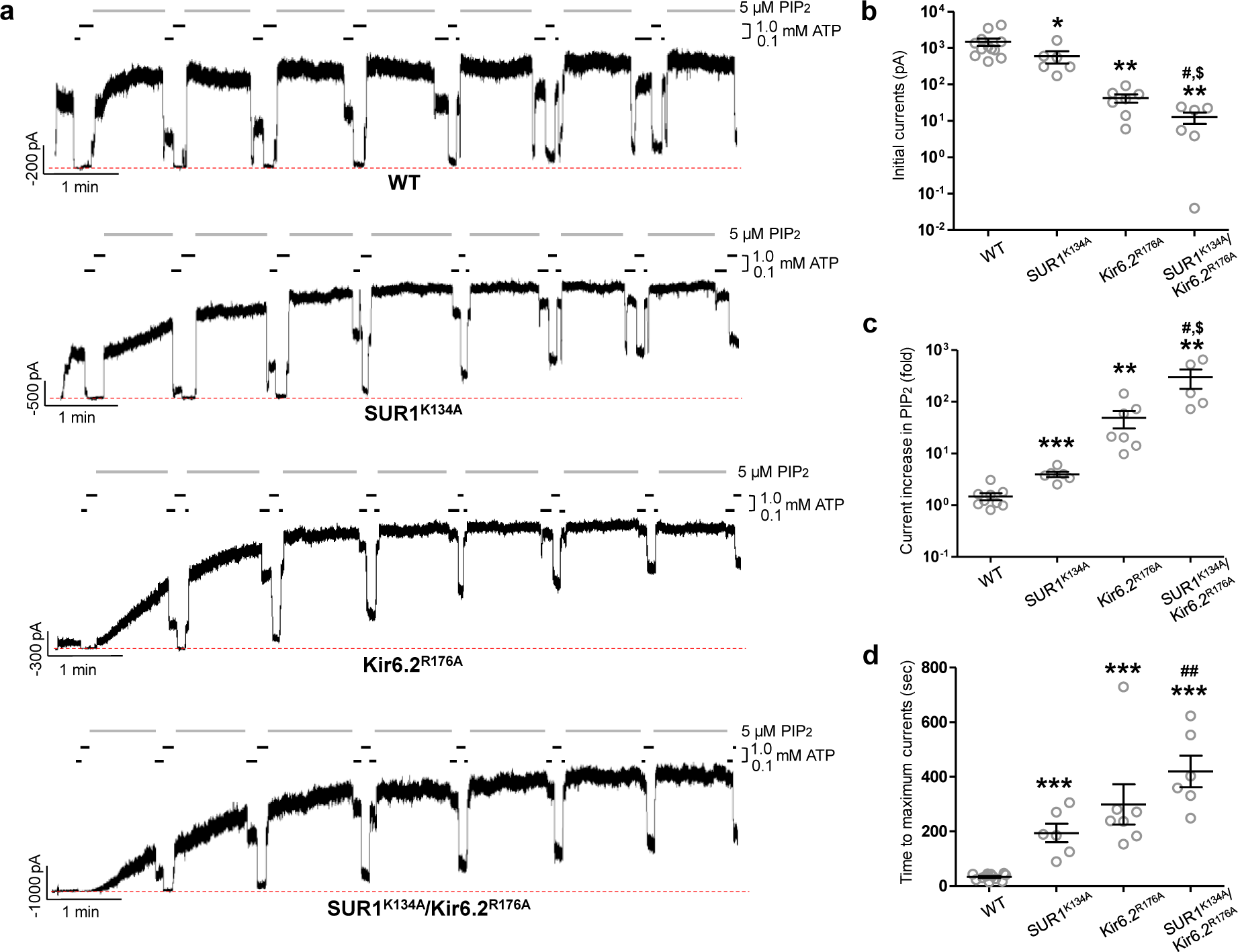
Functional testing of the two PIP_2_ binding sites. **(a)** Representative inside-out patch-clamp recordings (−50mV, inward currents shown as upward deflections; the red dashed line indicates 0 currents) of various channels: WT, SUR1^K134A^ (the second PIP_2_ binding site mutation)/Kir6.2, SUR1/Kir6.2^R176A^ (the first PIP_2_ binding site mutation), and SUR1^K134A^/Kir6.2^R176A^ double mutant, exposed alternately in 5 μM PIP_2_, 0.1 mM ATP, and 1 mM ATP as indicated by the bars above the recordings. The brief exposures to 0.1 mM and 1 mM ATP between PIP_2_ exposures were used to monitor the gradual decrease in ATP sensitivity as channel opening became increasingly stabilized by PIP_2_. **(b)** Group data of recordings shown in (a) comparing initial currents in K-INT at the time of patch excision. For each group, individual data points of 12 (WT), 6 (SUR1^K134A^), 7 (Kir6.2^R176A^), and 6 (SUR1^K134A^/Kir6.2^R176A^) patches and the mean ± SEM of the data points are shown. **(c)** Comparison of the maximum fold-increase in currents after PIP_2_ exposure in different channels from the same recordings analyzed in (b). **(d)** Comparison of the time of exposure in PIP_2_ for currents to reach maximum in different channels using the same recordings analyzed in (b) and (c). Note for (b) and (c), the y-axis is in log scale for better visualization. Statistical significance is marked by \**p*<0.05, \*\**p*<0.005, \*\*\**p*<0.001 between WT and the mutants, ^#^*p*<0.05, ^##^*p*<0.005 between SUR1^K134A^ and the double mutant, and ^$^*p*<0.05 between Kir6.2^R176A^ and the double mutant, using one-way ANOVA and Tukey’s post-hoc test.

### Kir6.2 pore in open conformation

In Kir channels, there are three constriction points in the K^+^ conduction pathway: the selectivity filter on the extracellular side, the inner helix gate at the helix bundle crossing where four inner helices (M2) converge, and the G-loop gate in the cytoplasmic pore just below the membrane^13^. The structure of the PIP_2_-bound SUR1/Kir6.2^Q52R^ channel has an open Kir6.2 inner helix gate (Fig. 4a). Clear cryoEM density shows rotation of the Kir6.2-F168 side chains away from the pore’s center, thus causing a dilated pore size (Figs. 4a, 5a, S4a, movie 1). The radius of the K^+^ pathway at the inner helix gate is ∼3.3 Å, compared to ∼0.5 Å in the WT K_ATP_ channel bound to inhibitors we reported previously^38,40^, exemplified by the ATP and repaglinide (RPG) bound structure (PDB ID 7TYS)^38^ (Fig. 4b,c). The radius at the inner helix gate in our current structure is comparable to that in the open structure of SUR1/Kir6.2^C166S,G334D^ (3.3 Å)^28^, and the pre-open structure of SUR1-Kir6.2^H175K^ fusion channel (3.0 Å)^29^ (Fig. S4b). In contrast to the inner helix gate, little difference is observed in the G-loop gate between closed and open structures (Fig. 4), suggesting minimum involvement of the G-loop gate in K_ATP_ channel gating by PIP_2_.

**Figure 4.**
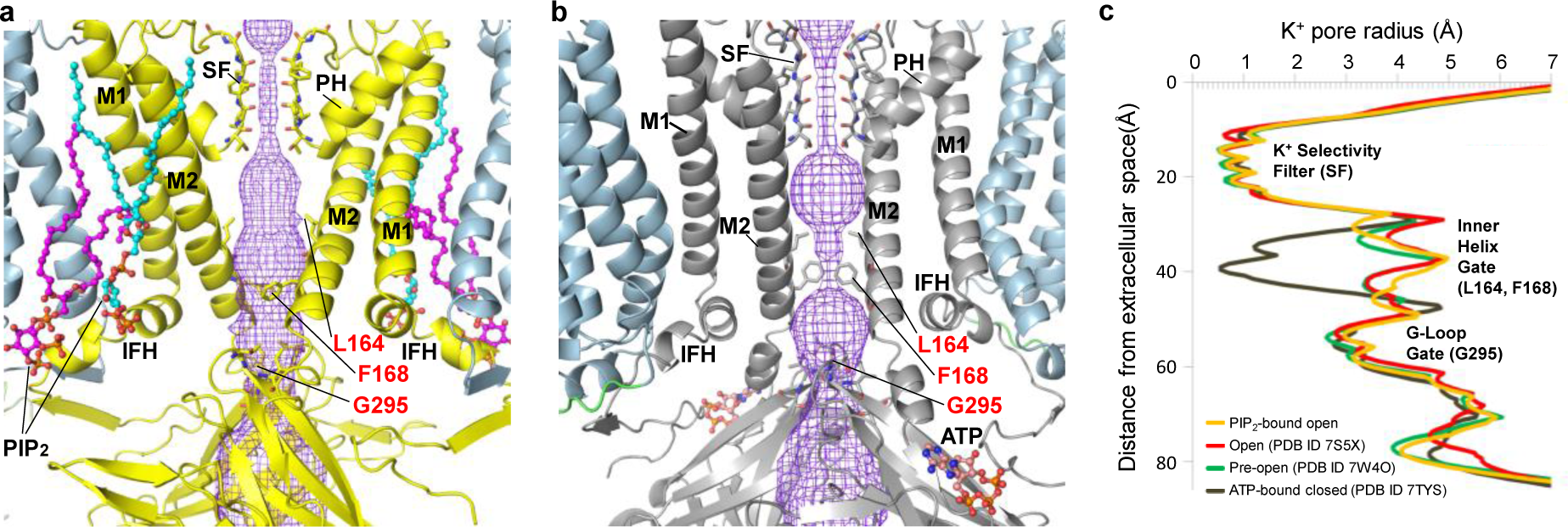
Comparison of the inner helix gate in the open and closed K_ATP_ channel structures. **(a)** Structure of Kir6.2^Q52R^ in a PIP_2_-bound open state. Bound PIP_2_ are shown as cyan and magenta sticks and spheres, and the side chains of inner helix gating residues L164 and F168 at the helix bundle crossing and G295 at the G-loop are shown as yellow sticks and labeled in red. The pore for the ion pathway (purple mesh) is constricted at the selectivity filter (SF) but open at the inner helix gate (3.3 Å). The transmembrane α-helices (M1 and M2), the interfacial α-helices (IFH) and the pore α-helices (PH) are labeled. **(b)** Kir6.2 in an ATP-bound closed state (PDB ID 6BAA) with similar labels as panel A for comparison. **(c)** Pore radii for ion conduction pathway plotted against the distance from the extracellular opening, with the location of the selectivity filter (residues 130-133), the inner helix gate (residues L164 and F168), and the G-loop gate (residues 294-297) shown. The pore radii were calculated using the program HOLE implemented in coot and viewed using a MOLE rendering in PyMOL.

**Figure 5.**
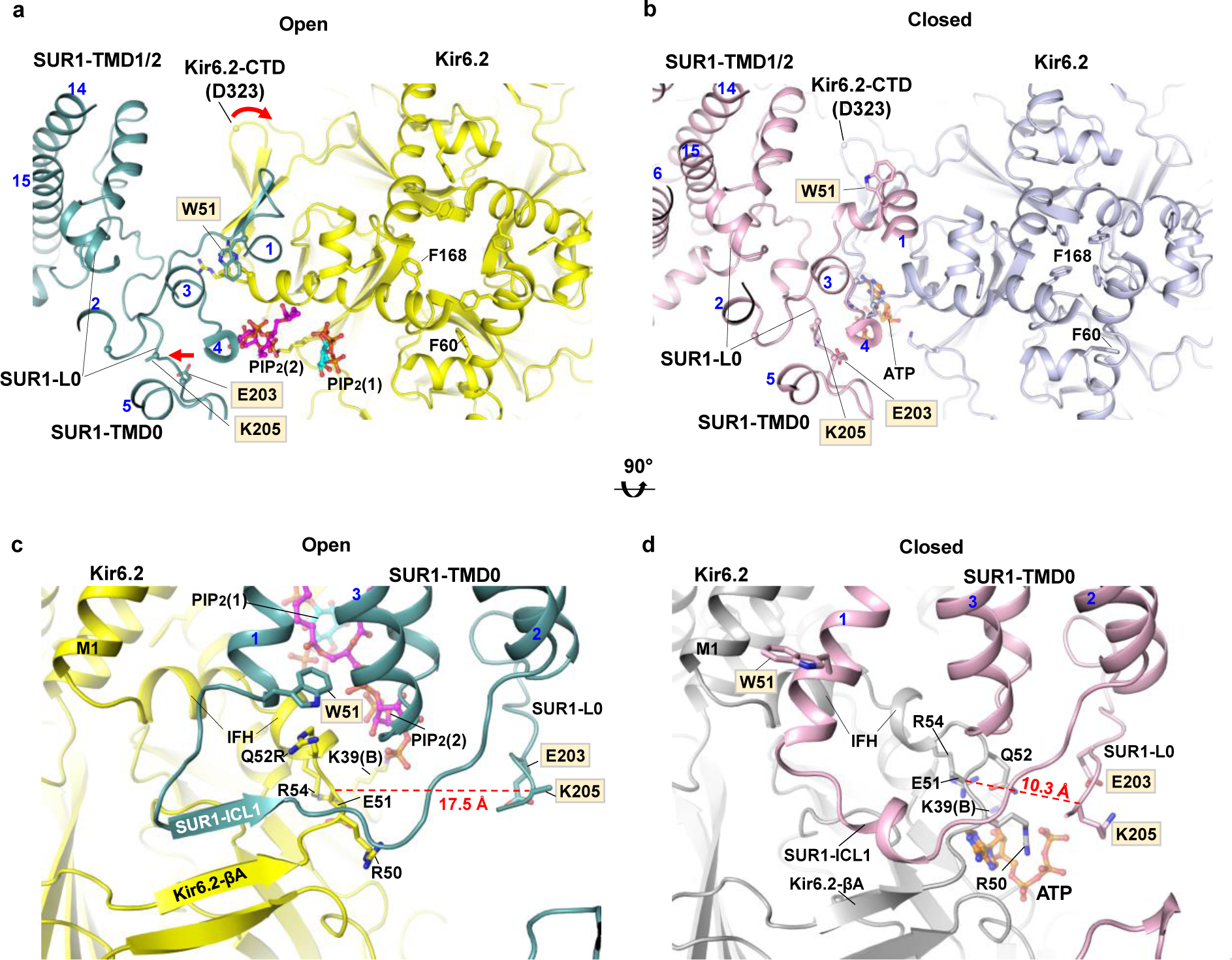
SUR1 and Kir6.2 cytoplasm-plasma membrane interface in open and closed K_ATP_ channel conformations. **(a)** SUR1-Kir6.2 cytoplasm-plasma membrane interface in open PIP_2_ (cyan and magenta sticks)-bound SUR1/Kir6.2^Q52R^ (SUR1 in teal and Kir6.2 in yellow) structure, and **(b)** closed ATP-(orange carbon sticks) and repaglinide (out of view)-bound WT (SUR1 in pink and Kir6.2 in grey) K_ATP_ channel (PDB ID 7TYS) viewed from the extracellular side. Note reorientation of side chains of the Kir6.2 inner helix gate residue F168 and M1 residue F60, as well as W51 at the bottom of TM1 of SUR1-TMD0 in the two conformations. In (a), a 6.4° clockwise rotation of the Kir6.2-CTD comparing the open to the closed conformation is indicated by the red curved arrow (with D323 Cα in each structure marked as spheres), and SUR1-L0, marked by the ATP-binding residue K205 and an adjacent residue E203, movement away from the ATP binding pocket in the open conformation relative to the closed conformation is marked by a red arrow. **(c)** A side view of the interface in the open conformation. SUR1-intracellular loop 1 (ICL1) forms a β-sheet with Kir6.2-βA just before the interfacial helix (IFH). Remodeling of residues involved in ATP binding and gating compared to the closed conformation in (d), including Kir6.2-R50, R54, E51 and K39(B) (“B” denotes the adjacent Kir6.2 subunit) as well as SUR1-E203 and K205 in L0 are shown. Most notably, the Kir6.2 mutant residue Q52R has its side chain reoriented to interact with the side chain of SUR1-W51 which also has its side chain reoriented from the closed conformation. Color scheme same as (a). **(d)** The same side view as in (b) but of the closed conformation. In all panels, SUR1 residue labels are outlined in grey. The red dashed line in (c) and (d) is the distance between the Cα of Kir6.2 E51 and SUR1 K205. In all panels, SUR1 helices are indicated with blue numbers.

The SUR1/Kir6.2^Q52R^ K_ATP_ sample was purified in the absence of nucleotides; as expected, no ATP cryoEM density was observed at the inhibitory ATP-binding site on Kir6.2. Nonetheless, the open conformation includes structural rearrangements at the ATP binding site that would disfavor ATP binding (Fig. 5a,c, S4a,c). Specifically, the sidechains of Kir6.2-K39 and R50 flip away from ATP-interacting positions, with the K39 sidechain now coordinating PIP_2_ at the second site; moreover, the sidechain of Kir6.2-E51 forms a salt bridge with R54 to partially occlude the ATP binding pocket (Fig. 5c). Our observation agrees with previous hypotheses that the open channel conformation is not compatible with ATP binding at Kir6.2 and that ATP inhibits the channel by stabilizing the channel in the closed conformation^16^.

Relative to channels bound to inhibitors, in which the Kir6.2-CTD is predominantly positioned close to the plasma membrane (“CTD-up” conformation)^38^, the SUR1/Kir6.2^Q52R^ open channel structure shows the Kir6.2-CTD is also juxtaposing the membrane but further rotated clockwise (viewed from the extracellular side) (compare Fig. 5a and 5b, movie 1). This twisting as well as the other structural changes at the ATP binding site discussed above are similarly observed in the open structure of the SUR1/Kir6.2^C166S,G334D^ channel^28^, and the pre-open structure of the SUR1-Kir6.2^H175K^ fusion channel^29^ (Fig. S4b,d). The converging observations in three open structures using different mutant constructs suggest WT K_ATP_ channels undergo similar conformational changes between closed and open states.

### Analysis of SUR1 structure

In the PIP_2_-bound open SUR1/Kir6.2^Q52R^ structure, the two NBDs of SUR1 are not well resolved; in particular, NBD2 is highly dynamic with a local resolution > 5 Å (Figs. 1, S1d). However, the two NBDs are clearly separated, as predicted since no MgATP or MgADP was added to the sample. Notably, no cryoEM density corresponding to the distal N-terminal peptide of Kir6.2^Q5 2^ ^R^ was observed in the central cleft lined by TM helices from TMD1 and TMD2 of the SUR1-ABC core even when the map was filtered to 7 Å (Fig. S5a); no continuous density corresponding to KNtp was observed at an alternative site to suggest a stable resting position, precluding structural modeling of this flexible domain. The absence of KNtp cryoEM density in the PIP_2_ - bound open SUR1/Kir6.2^Q52R^ structure is in contrast to the clear KNtp density observed in the central cleft of SUR1-ABC core in the apo closed SUR1/Kir6.2^Q52R^ structure at comparable resolution and contour level (Fig. S2e, S5). Presence of KNtp in the SUR1-ABC core has been previously reported in inhibitor-bound structures and linked to channel closure^38,41,42^. Worth noting, in a publsihed open K_ATP_ structure where NBDs are bound to MgATP/MgADP and dimerized, KNtp density was also absent^28^. The lack of KNtp in the SUR1-ABC core in the open SUR1/Kir6.2^Q52R^ structure further enforce the correlation between the absence of KNtp in the central cleft of the SUR1-ABC core and channel opening, whether SUR1-NBDs are dimerized or not.

The ABC modules of the four SUR1 subunits in the SUR1/Kir6.2^Q52R^ map from a C4 non-uniform refinement adopt a propeller-like conformation (Fig. 1a, b). Further 3D classification of the symmetry expanded particle set revealed three distinct rotational positions of SUR1-ABC module relative to the Kir6.2-SUR1-TMD0 tetrameric core (Fig. S6a,b). Similar conformational dynamics have been reported for the SUR1 NBDs-dimerized, open structure of the SUR1/Kir6.2^C166S,G334D^ channel^28^, thus likely represent intrinsic flexibility of this domain regardless of nucleotide binding and/or NBD dimerization. In addition to the dynamics of the SUR1-ABC module within the PIP_2_-bound open SUR1/Kir6.2^Q52R^ conformation class, more pronounced heterogeneity was also captured in 3D classification of symmetry expanded particles that include all particles from 2D classes (Fig. S2). Here, rotation of the SUR1-ABC module is seen correlated with the corkscrew position of the Kir6.2-CTD, with the SUR1-ABC module rotated in towards the Kir6.2 tetramer in the apo closed CTD-down conformation class, and more flung out in the PIP_2_-bound open CTD-up conformation (Figs. S2c,d, 6c,d). Such correlated dynamic movements of SUR1 and Kir6.2 may reflect the conformational transition between closed and open channels.

**Figure 6.**
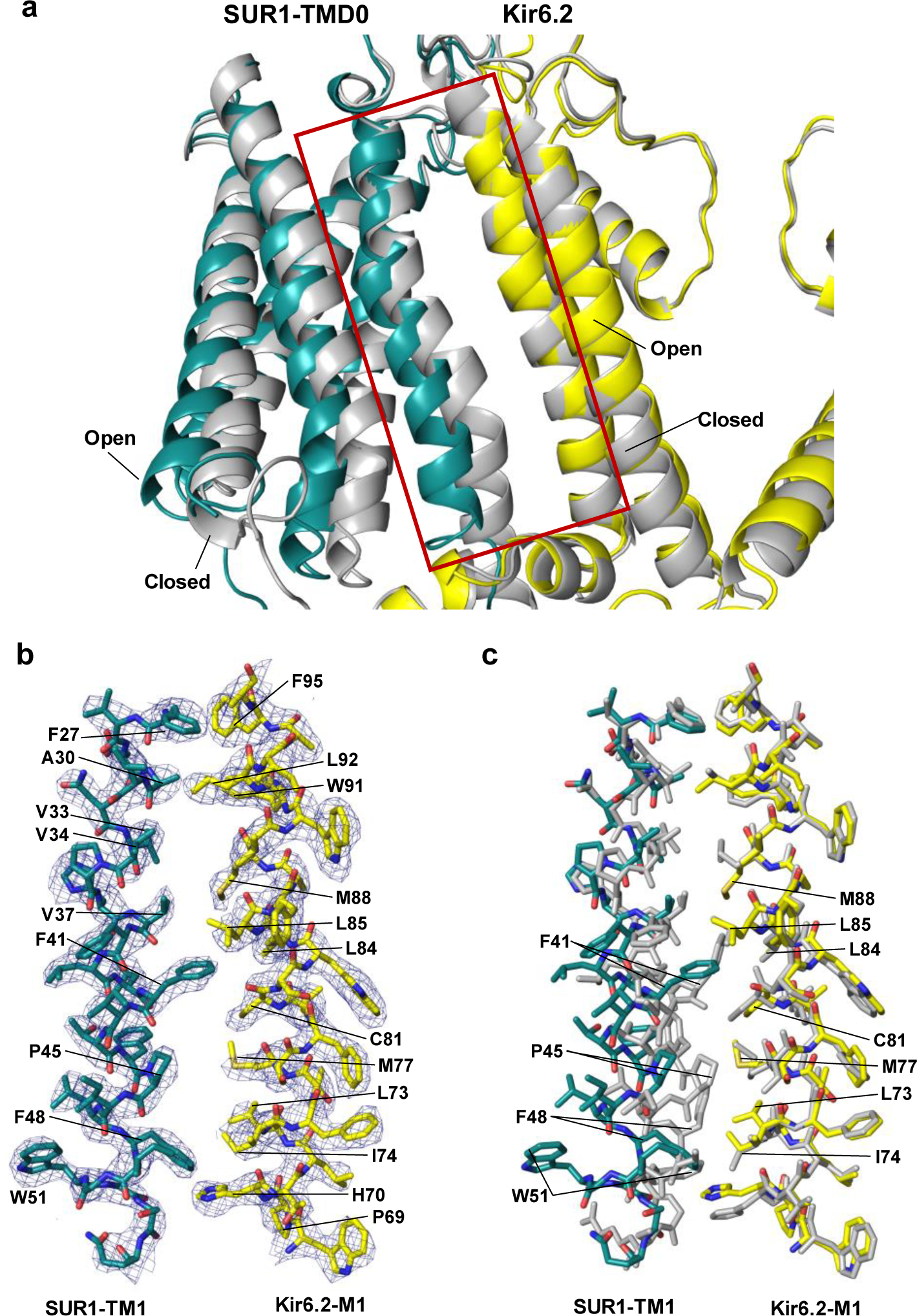
Structural changes between open- and closed-forms of the K_ATP_ channel at the SUR1 and Kir6.2 TM interface. **(a)** An overlay of the SUR1-Kir6.2 TM interface between open SUR1/Kir6.2^Q52R^ K_ATP_ channel and the closed K_ATP_ channel (PDB ID 6BAA) showing the outward movement of the cytoplasmic half of the SUR1-TMD0. In the open structure, SUR1 is shown in teal and Kir6.2 in yellow. The closed structure is shown in grey. **(b)** Close-up view of the open structure in the boxed region in (a). CryoEM map (blue mesh, 0.17V/8.5σ map contour) is superposed with the structural model. For clarity, PIP_2_ is omitted. **(c)** An overlay of the open with the closed K_ATP_ channel structures in the boxed region in (A) (same color scheme) showing significant changes in residues of SUR1-TM1 at the inner leaflet half of the plasma membrane.

Compared to inhibitor-bound closed Kir6.2/SUR1 channel structures [reviewed in^6,7^], SUR1 subunits in the SUR1/Kir6.2^Q52R^ open structure are more tilted away from Kir6.2 and elevated towards the plasma membrane (viewed from the side) such that they are nearly in plane with the Kir6.2 tetramer (compare Fig. 1b and 1c). Accompanying this upward SUR1 motion, there is an overall rigid-body rotation of SUR1 driven by a rotation of the TMD0 of SUR1 in the open structure relative to closed structures (Fig. 6). At the outer leaflet, SUR1-F41, which interacts with Kir6.2-L84 in the closed conformation, now interacts with Kir6.2-L85 in the open conformation (Fig. 6c). At the inner leaflet, TMD0 moves away from Kir6.2 (Fig. 6a, 6c), creating a space between SUR1-TM1 and Kir6.2-M1. Using PISA analysis^43^, the contact surface area at this interface (between SUR1-TM1 residues 27-54 and Kir6.2-M1 residues 66-96) is 645.6 Å^2^ (ΔG = −20 kcal/mol) in the closed channel structure (PDB ID 6BAA), which is reduced to 395.9 Å^2^ (ΔG = −12 kcal/mol) in the SUR1/Kir6.2^Q52R^ open structure. Binding of a second PIP_2_ in this space (see Fig. 2b) compensates for the lost surface contact between SUR1-TMD0 and Kir 6.2, thus stabilizing this interface in the open channel.

Similar structural changes were also described in the pre-open SUR1-Kir6.2^H175K^ fusion structure where SUR1 NBDs are bound to MgATP/MgADP and dimerized^30^ (Fig. S4). Dimerization of SUR1 NBDs antagonizes ATP inhibition at Kir6.2 and stimulates K_ATP_ channel activity. It has been proposed that NBD dimerization causes outward bending of the cytoplasmic half of SUR1-TMD0 and the pulling away of K205 in SUR1-L0 from binding ATP at the inhibitory site to stimulate channel activity^30^. Since our open SUR1/Kir6.2^Q52R^ structure does not contain MgATP/MgADP and the NBDs are clearly separated, the similar conformational changes we observed at SUR1-TMD0/L0 are not driven by SUR1 NBD dimerization. One possibility is that MgATP/MgADP and PIP_2_ converge on the same structural mechanism to open the channel, as has been proposed previously based on functional studies^44^.

### Kir6.2-Q52R stabilizes the open conformation by interacting with SUR1-W51

Kir6.2-Q52 is located immediately N-terminal to the interfacial helix (IFH; a.k.a. slide helix) and C-terminal to βA (Fig. 5c,d). In published ATP-bound closed structures, Kir6.2-Q52 lies close to SUR1-E203 in the proximal portion of the SUR1-L0 linker (Fig. 5d). Previous studies have shown that enforcing Kir6.2-Q52 and SUR1-E203 interactions via engineered charged amino acid pair or cysteine crosslinking reduced ATP inhibition IC_50_ by 100-fold or caused spontaneous channel closure in the absence of ATP, respectively^45^. These results indicate that stabilizing the interface between Kir6.2 βA-IFH and SUR1-L0 stabilizes ATP binding and channel closure. In contrast, in the SUR1/Kir6.2^Q52R^ open structure, the SUR1-L0 linker moves ∼7 Å towards SUR1 and away from the ATP-binding site on Kir6.2 (compare Fig. 5c and 5d), with SUR1-E203 and K205 flipped away from Kir6.2 such that SUR1-TMD0 ICL1 (aa 52-60) engages with Kir 6.2 - βA to form a main chain β-sheet (Fig. 5c). In particular, the sidechain of Kir6.2 residue 52 (Q in WT, R in mutant) now faces SUR1-TMD0 and interacts with SUR1-W51 (Figs. 5c, S4f), tethering the Kir6.2-CTD to SUR1-TMD0 in a rotated open position.

The Kir6.2-Q52R mutation increases *P_o_* and decreases ATP inhibition of K_ATP_ channels in a SUR1-dependent manner^46^. The cation-π interaction observed between Kir 6.2-Q52R and SUR1-W51 in the open conformation (Fig. 7a) led us to hypothesize this interaction underlies the gain-of-function gating effect of the Kir6.2-Q52R mutation, by stabilizing Kir6.2-CTD in a rotated open conformation. To test this, we mutated W51 of SUR1 to cysteine and assessed the ATP sensitivity of channels formed by co-expressing Kir6.2^Q52R^ and SUR1^W51C^ using inside-out patch-clamp recording. SUR1^W51C^ reversed the effect of Kir6.2^Q52R^ such that the ATP sensitivity of the channel resembles WT channels (Fig. 7b). The IC_50_ values of ATP inhibition for WT, SUR1/Kir6.2^Q52R^, SUR1^W51C^/Kir6.2, and SUR1^W51C^/Kir6.2^Q52R^ channels are 6.25 ± 0.44 μM, 161.2 ± 16.23 μM, 20.88 ± 1.96 μM, and 8.98 ± 0.49 μM, respectively (Fig. 7c). Corroborating these findings, in Rb^+^ efflux assays cells co-expressing Kir6.2^Q52R^ and SUR1^W51C^ showed efflux levels similar to cells expressing WT channels, in contrast to the significantly higher efflux in cells co-expressing Kir6.2^Q52R^ and WT-SUR1 (Fig. 7d). These results provide strong evidence that Kir6.2-Q52R interacts with SUR1-W51 to enhance channel activity and reduce ATP inhibition, thus explaining the SUR1-dependent pathophysiology of this neonatal diabetes mutation.

**Figure 7.**
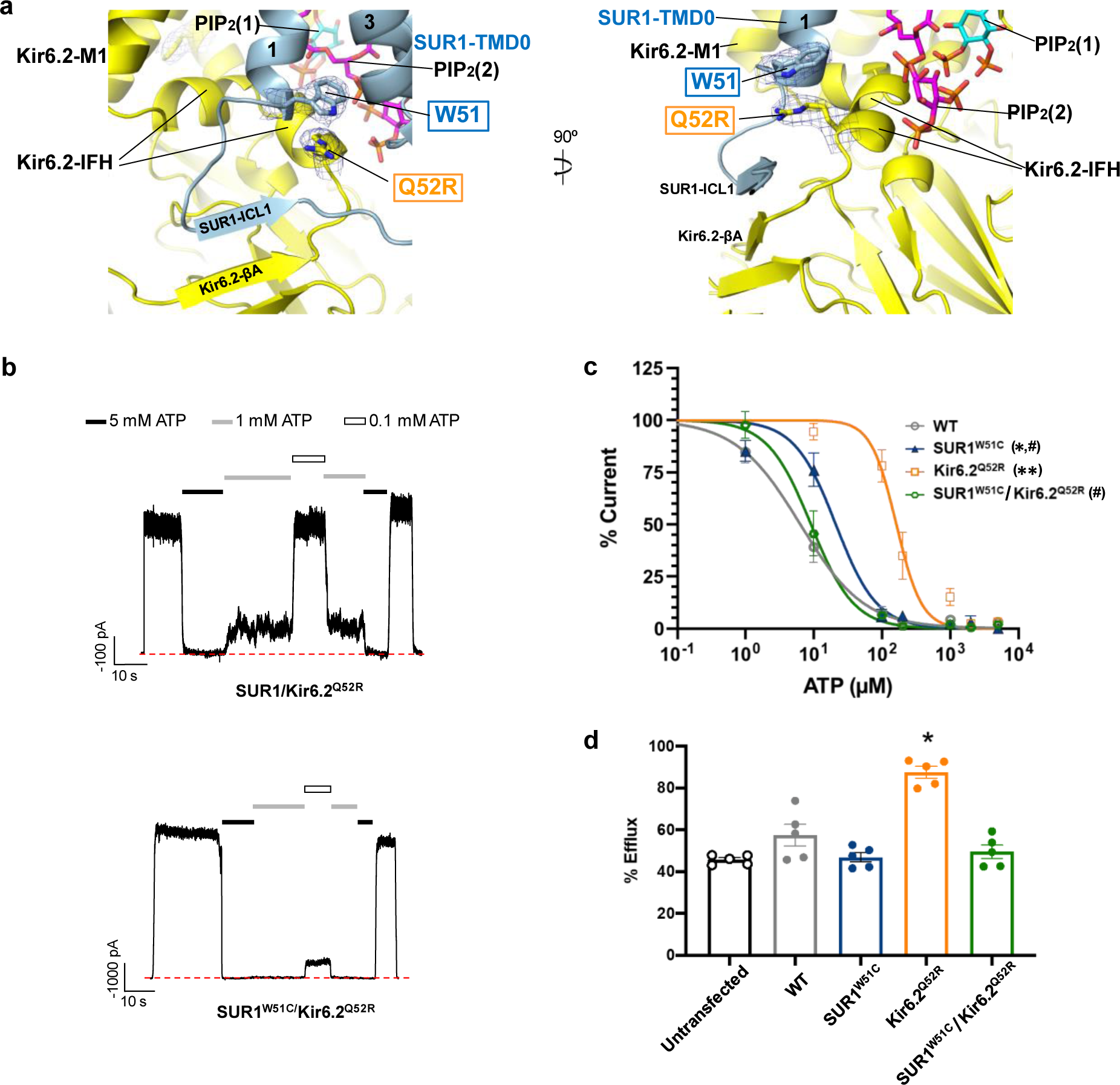
Kir6.2-Q52R interacts with SUR1-W51 to reduce channel sensitivity to ATP inhibition and enhance channel activity. **(a)** A close-up view of the interaction between Kir6.2-Q52R and SUR1-W51 with cryoEM map (blue mesh, 0.17V/8.5σ contour) superposed with the structural model in two different angles. **(b)** Representative inside-out patch-clamp recordings (−50mV, inward currents shown as upward deflections; red dashed line indicates baseline 0 currents) of the SUR1/Kir6.2^Q52R^ channel and SUR1^W51C^/Kir6.2^Q52R^ channel exposed to differing concentrations of ATP as indicated by the bars above the recordings. **(c)** ATP dose response of WT (SUR1/Kir6.2) channels, channels containing the SUR1^W51C^mutation (SUR1^W51C^), Kir6.2^Q52R^ mutation (Kir6.2^Q52R^), or both SUR1^W51C^ and Kir6.2^Q52R^ mutations (SUR1^W51C^/Kir6.2^Q52R^). Curves were obtained by fitting the data points (mean ± SEM of 3-8 patches) to the Hill equation (see Methods). \**p* < 0.01, \*\**p* < 0.0001 compared to WT; #*p* < 0.0001 compared to Kir6.2^Q52R^, one-way ANOVA with Tukey’s *post hoc* test **(d)** Rb^+^ efflux of various channels (same labeling as in (c)) expressed in COSm6 cells. Untransfected cells were included to show background efflux. Only SUR1+Kir6.2^Q52R^ channels showed significant activity above WT (*p* < 0.0001, n=5; one-way ANOVA with Tukey’s *post hoc* test).

To test whether the interaction between Kir6.2-Q52R and SUR1-W51 that gives r ise to the enhanced activity and reduced ATP sensitivity is dependent on PIP_2_ binding at the two sites revealed by our structure, we tested the effect of Kir6.2^Q52R^ on the background of SUR1^K134A^, Kir6.2^R176A^, or SUR1^K134A^ and Kir6.2^R176A^, to weaken the novel PIP_2_ binding site, the conserved PIP_2_ binding site, or both. Inside-out patch-clamp recording experiments showed a leftward shift in the dose response of ATP inhibition of the SUR1^K134A^/Kir6.2^Q52R^ channel (IC_50_= 54.08 ± 7.14 μM, H = 1.55), the SUR1/Kir6.2^Q52R/R176A^ channel (IC_50_=17.97 ± 1.20 μM, H = 1.41), and the SUR1K^134A^/Kir6.2^Q52R/R176A^ channel (IC_50_= 8.30 ± 1.11 μM, H = 1.18) relative to the SUR1/Kir6.2^Q52R^ channel (IC_50_= 161.20 ± 16.23 μM, H = 2.34) (Fig. S7a). SUR1^K134A^ or Kir6.2^R176A^ alone had little effect on the IC_50_ of ATP inhibition (4.12 ± 0.30 μM, H = 1.54, and 11.27 ± 1.95 μM, H = 0.84, respectively, compared to 6.25 ± 0.44 μM, H = 0.93 for WT channels). In agreement, Rb^+^ efflux experiments showed attenuation of the gain-of-function phenotype associated with Kir6.2^Q52R^ when combined with PIP_2_ binding mutations SUR1^K134A^, Kir6.2^R176A^, or both (Fig. S7b). Taken together, these results support a role of PIP_2_ at the two binding sites in manifesting the gain-of-function effect of Kir6.2^Q52R^.

## Discussion

In this study, we surmounted the technical difficulties in capturing PIP_2_ bound to K_ATP_ channel, firstly by using a neonatal-diabetes Kir6.2^Q52R^ variant that enhances channel activity in an SUR1-dependent mechanism, and secondly by using natural brain-derived long-chain PIP_2_ before membrane solubilization, rather than the previously used approach of adding short-chain synthetic diC8-PIP_2_ after channel purification^24,30,47^. The cryoEM map of the lipid densities with long acyl chain tails, together with functional studies on designed K_ATP_ channel mutants, permitted the assignment of PIP_2_ at both the expected site and the novel binding site (Figs. 2b,c, S3b, S4a,b). The novel PIP_2_ binding site, coordinated by Kir6.2 and its regulatory subunit SUR1, uncovers an unprecedented cooperation between an ion channel and transporter on PIP _2_ binding, and highlights the uniqueness of the K_ATP_ complex in the evolution of K^+^ channels and ABC proteins. The structure resolves a major problem in the long-standing puzzle of how SUR1, in particular TMD0 of SUR1, modulates the open probability of the Kir6.2 channel^25,26^. Moreover, it answers a long-standing question of how a prominent clinical Kir6.2 mutation increase s channel activity to cause neonatal diabetes^46^.

### Unique PIP_2_ binding pocket in K_ATP_ channels

As a member of the Kir channel family, Kir6.2 has a homologous PIP2 binding site akin to that identified in Kir2 and Kir3^17–20^. Surprisingly, our structure reveals a second, novel PIP _2_ binding site immediately next to the conserved Kir channel PIP_2_ binding site. This second site is uniquely coordinated by both SUR1 and Kir6.2 subunits. Mutational studies support functional importance of PIP_2_ binding at both sites (Fig. 3). In single channel recordings, Kir6.2 channels lacking SUR1 have low *P_o_* with brief openings, in contrast to Kir6.2 channels assembled with SUR1 or SUR1-TMD0, which exhibit ∼10-fold higher *P_o_* and long bursts of openings^16,25,26,48^. We propose the low *P_o_* and brief openings in Kir6.2 channels lacking SUR1 result from PIP_2_ binding at the conserved site, which tethers Kir6.2-CTD near the membrane to rotate to the open position. PIP_2_ binding at this site may not be as strong as in Kir2 and Kir3 channels due to sequence variations at key residues including Kir6.2 P69 and H175 (both positively charged amino acids in Kir2 and 3)^17–20^, explaining the low *P_o_* of Kir6.2. PIP_2_ binding at the unique second site formed by SUR1 and Kir6.2 enables SUR1 to stabilize Kir6.2-CTD in the rotated open position, giving rise to bursts of openings and higher *P_o_*.

The PIP_2_ binding pocket is smaller in the previously published inhibitor-bound closed structure than in the SUR1/Kir6.2^Q52R^ open structure. In a solvent accessible surface representation of the model for a closed structure (exemplified by the repaglinide/ATP bound PDB ID 7TYS)^38^, PIP_2_ at the second site would clash with SUR1 (Fig. S8b). Inspection of the closed SUR1/Kir6.2^Q52R^ CTD-down conformation (see Fig. S2) also finds that PIP_2_ binding at the second site would clash with SUR1 (Fig. S8c), indicating this structure represents an apo state, at least with respect to PIP_2_ binding at the second novel site. These observations imply that PIP_2_ moves in and out of the second site as the channel opens and closes. In this regard, it is interestingly to note that a conformation-dependent change in the PIP_2_ binding site in the voltage-dependent KCNQ1 (Kv7.1) channel was recently reported, where movement of the voltage sensing transmembrane helix S4 up or down the membrane is associated with opening or occlusion of the PIP_2_ binding site to open or close the channel, respectively^49^. A question that arises is whether smaller lipids may occupy the more constricted second site as the channel transitions from the open to the closed conformation. Due to resolution differences, it is not possible to resolve lipid density in the apo closed SUR1/Kir6.2 ^Q52R^ channel map. However, in our published closed structures without exogenous PIP_2_, there was also lipid density, which we tentatively modeled as two phosphatidylserines^38^. Another question is whether the first PIP_2_ binding site is always occupied by PIP_2_ since it appears available in both the open and closed channel conformations (Fig. S8). More studies are needed to answer these questions.

K_ATP_ channels exhibit lower specificity towards PIPs compared to other Kir channels, and are activated equally well by PI(4,5)P_2_, PI(3,4)P_2_, and PI(3,4,5)P_3_, and also by PI(4) P and long chain (LC)-CoAs^33^. The structural basis underlying the reduced PIPs specificity for K_ATP_ channel activation requires further investigation. The aforementioned sequence variations at key PIP_2_ coordinating residues at the site homologous to other Kir channels may contribute to the lower specificity. Additionally, the large size of the tandem PIP_2_ binding pocket may accommodate different PIPs and LC-CoAs. Such degeneracy may account for the observation that purified K_ATP_ channels reconstituted in lipid bilayers lacking PIP_2_ exhibited spontaneous single channel openings^28^, in contrast to Kir2 and Kir3 channels which require PIP_2_ for activity^20,50^.

### Mechanism of PIP_2_ and ATP antagonism

PIP_2_ and ATP functionally compete to open or close K_ATP_ channels (Fig. 3a)^11,12^. Kinetic analyses have indicated that PIP_2_ and ATP binding are mutually excluded^16^. Comparison between PIP_2_-bound SUR1/Kir6.2^Q52R^ open structure and previously published ATP-bound closed structures shows PIP_2_ and ATP antagonism occurs via both binding competition and allosteric mechanisms (Fig. 5, movie 1). At the level of binding competition, both ligands compete for a common binding residue, Kir6.2-K39, which interacts with ATP in the ATP-bound closed structures^38,40^ but with PIP_2_ in the PIP_2_-bound open structure (Figs. 2b, 5c). Previous molecular dynamics simulations, using ATP-bound closed structures of Kir6.2 tetramer^51^ or tetramer of Kir6.2 plus SUR1-TMD0^38^ with a single PIP_2_ molecule docked in the conserved PIP_2_ binding site, found that Kir6.2-K39^51^ or both K39 and R54^38^ switched between ATP binding and PIP_2_ binding. In the SUR1/Kir6.2^Q52R^ open structure, we see Kir6.2-K39, rather than binding PIP_2_ in the conserved site, binds the second PIP_2_ in the novel site. In contrast, Kir6.2-R54 is not involved in PIP_2_ binding; instead, it hydrogen bonds with Kir6.2-E51 in the same subunit and E179 in the adjacent Kir6.2 subunit, which stabilizes the interface between the IFH and the C-linker helix (the linker helix connecting M2 and C-terminal cytoplasmic domain of Kir 6.2) in the open conformation. Mutation of both K39 and R54 to alanine has been shown to reduce ATP as well as PIP_2_ sensitivities^52^. Our structure clarifies the structural role of K39 and R54 and suggests while K39A likely reduces PIP_2_ sensitivity by weakening PIP_2_ binding at the second site, R54A likely reduces PIP_2_ sensitivity indirectly by disrupting interactions in Kir6.2 that are needed to stabilize the open channel conformation.

Allosterically, PIP_2_ binding diverts ATP binding residues away from the ATP binding pocket to disfavor ATP binding. These include sidechain reorientation of Kir 6.2 K39 and R50, and interaction between Kir6.2 R54 and E51 that partially occludes the ATP binding pocket (Fig. 5c). Moreover, in the ATP-bound closed conformation, SUR1-L0 interfaces with Kir 6.2 - βA, allowing SUR1-K205 to coordinate ATP binding^38,42,45,53^ and stabilize channel closure. However, in the open structure, SUR1-L0 is disengaged from Kir6.2-βA, allowing Kir6.2-CTD to rotate such that ICL1 of SUR1-TMD0 engages with Kir6.2-βA, forming a continuous β-sheet to stabilize the open conformation, which also disfavors ATP binding. Similar changes have been described in published SUR1/Kir6.2^C166S,G334D^ open structure^28^, and SUR1-Kir6.2^H175K^ fusion pre-open structure^30^ (Fig. S4f). The common structural rearrangements from closed to open state seen independent of mutations support the same gating transition in WT channels and highlight a key role of SUR1 in stabilizing the Kir6.2-CTD in two distinct rotational positions to close or open the K_ATP_ channel.

#### Insights on disease mutations

The study provides mechanistic insight on how Kir6.2^Q52R^ causes neonatal diabetes. The strong cation-π interaction engendered by the Kir6.2-Q52R mutation with SUR1-W51 illustrates how changes at this interface have profound effects on channel gating and physiology. In the published SUR1/Kir6.2^G334D,C166S^ open structure and the SUR1-Kir6.2^H175K^ fusion channel pre-open structure Q52 of Kir6.2 is similarly in position to interact with W51 of SUR1^28,29^ (see Fig. S4f). The polar-π interaction between glutamine and tryptophan is much weaker than the cation-π interaction between arginine and tryptophan. The difference in binding energy for the relevant gas-phase interactions for polar-π is 1.4 kcal/mol (NH_3_-Benzene), compared to the cation-π interaction energy of 19.0 kcal/mol (NH_4_^+^-Benzene)^54,55^. That Q52 and Q52R in Kir 6.2 both interact with W51 of SUR1 suggests an important role of Kir6.2-Q52 in stabilizing Kir6.2-CTD and SUR1-TMD0 interface for channel opening. Functional studies showing that weakening PIP_2_ binding at either sites attenuate both WT and SUR1/Kir6.2^Q52R^ activity indicates endogenous PIP_2_ is required to support the activity of both WT and mutant channels. In the absence of normal PIP_2_ binding, the gain-of-function effect of Kir6.2^Q52R^ becomes much weaker. The presence of a minor conformation class corresponding to an apo closed SUR1/Kir 6.2 ^Q52R^ channel (Fig. S2) even in the presence of excess PIP_2_ further strengthens the notion that Kir6.2^Q52R^ alone is insufficient to cause disease in the absence of PIP_2_.

In addition to Kir6.2^Q52R^, many disease mutations are located at the SUR1-Kir6.2 interface seen in the SUR1/Kir6.2^Q52R^ open structure (Fig. 8). These include congenital hyperinsulinism (HI) associated loss-of-function Kir6.2 mutations: R54C, L56G, K67D, R176H and R177W, and SUR1 mutations: I46T, P133R, and L135V, as well as neonatal diabetes (ND) and <u>D</u>evelopmental delay, <u>E</u>pilepsy, and <u>N</u>eonatal <u>D</u>iabetes (DEND) syndrome associated gain-of-function Kir6.2 mutations: K39R, E51A/G, Q52L/R. G53D/N/R/S/V, V59A/G/M, W68C/G/L/R, K170N/R/T, E179A/K, and SUR1 mutations: P45L, I49F, F132L/V, and L135P (Fig. 8) ^56^. Some of these, such as HI-associated K67D and R176H, and ND/DEND associated K39R, W68C/G/L/R involve residues that coordinate PIP_2_ binding; others however, likely affect PIP _2_ or ATP gating allosterically by stabilizing channels in closed or open conformations.

**Figure 8.**
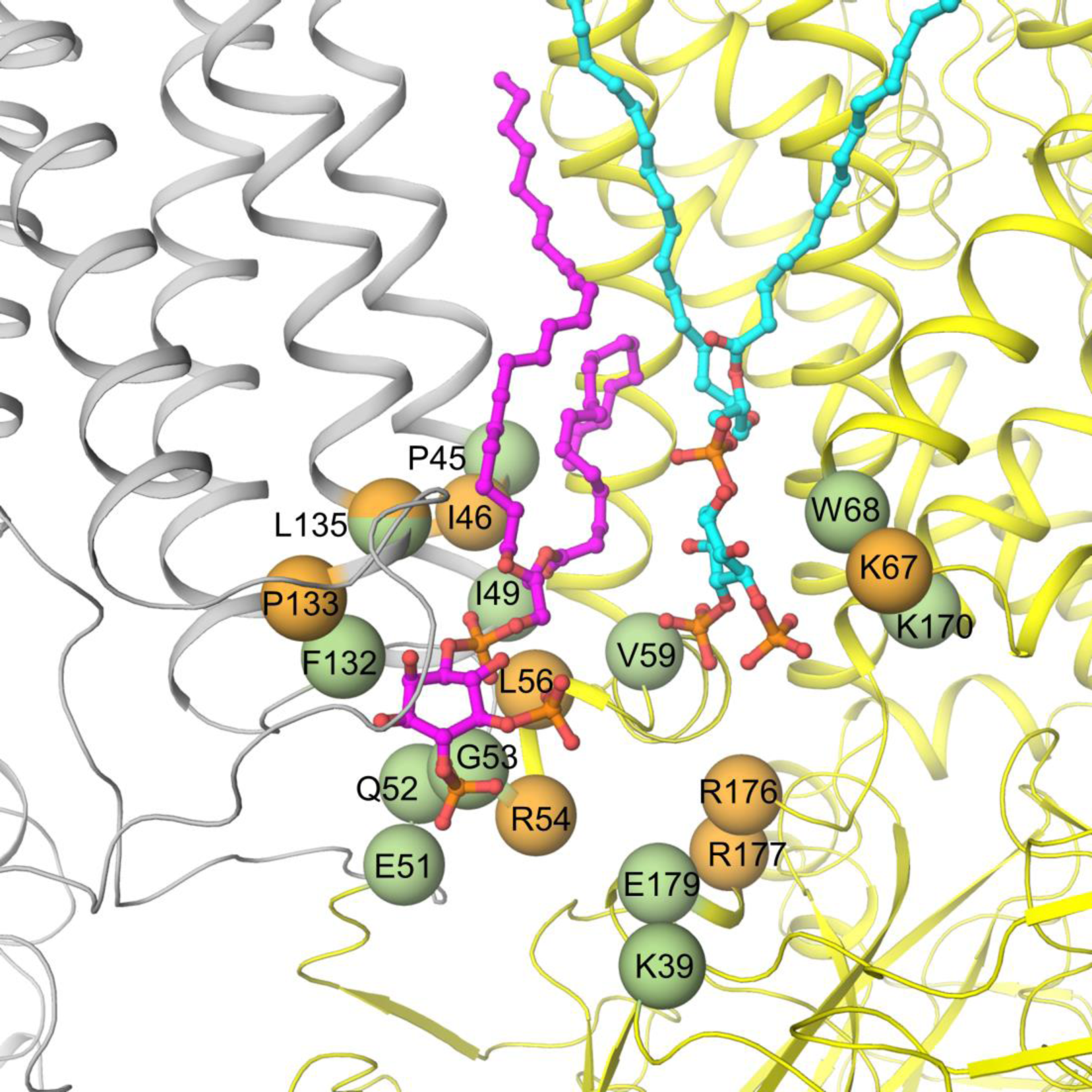
K_ATP_ channel disease mutations near the PIP_2_ binding pocket. Residues with variants that have been reported to cause neonatal diabetes/DEND syndrome are shown as green spheres and congenital hyperinsulinism shown as orange spheres (with the exception of L135, which is colored half green and half orange as mutation at this position has been linked to both diseases depending on the substituting amino acid). Neonatal diabetes/DEND syndrome mutations include SUR1-P45L, I49F, F132L/V, L135P, and Kir6.2-K39R, E51A/G, Q52L/R, G53D/N/R/S/R/V, V59A/G/M, W68C/G/L/R, K170N/R/T, E179A/K. Congenital hyperinsulinism mutations include SUR1-I46T, P133R and L135V, and Kir6.2-R54C, L56G, K67D, R176H, and R177W. SUR1 and Kir6.2 are colored grey and yellow, respectively.

In summary, the PIP_2_-bound K_ATP_ channel structure elucidates the intimate partnership between SUR1 and Kir6.2 in coordinating PIP_2_ binding and channel gating. The unique PIP_2_ binding pocket and the molecular interactions involved in gating provide a framework for designing new K_ATP_ modulators to control channel activity. The finding that SUR1 binds PIP_2_ to regulate Kir6.2 also raises the question whether PIP_2_ serves structural and functional roles in other ABC proteins, as has been implicated for CFTR^57^.

## Materials and methods

### Cell lines

COSm6 cells were cultured in high-glucose DMEM medium (GIBCO) supplemented with 10% Fetal Bovine Serum (Fisher Scientific), 100 U/mL penicillin, and 100 U/mL streptomycin at 37°C with 5% CO_2_.

### Protein expression and purification

Genes encoding rat Kir6.2^Q52R^ and N-terminal FLAG-tagged (DYKDDDDK) hamster SUR1 were cloned into pShuttle vectors and then the AdEasy vector (Stratagene), and packaged into recombinant adenoviruses, which were used for protein expression as described previously ^58^. However, instead of inducing K_ATP_ channel expression in INS-1 cells, which are pancreatic β cell derived and have endogenous WT Kir6.2 and SUR1 expression that would mix with the Q52R variant, a different mammalian adhesion cell line was used (COSm6 cells). COSm6 cells grown to mid-log in 15 cm tissue-culture plates were infected with adenoviruses packaged with Kir 6.2, SUR1, and tTA, using multiplicity of infections (MOIs) optimized empirically. Note the pShuttle vector used for SUR1 contains a tetracycline-regulated response element, necessitating co-infection of a tTA (tetracycline-controlled transactivator) adenovirus for SUR1 expression. At ∼24 hours post-infection, cell medium was changed and included 100 μM tolbutamide post-infection. Tolbutamide enhances K_ATP_ channel expression at the plasma membrane and washes out easily. At ∼48 hours post-infection, cells were harvested by scraping and cell pellets were frozen and stored at −80°C until purification.

SUR1/Kir6.2^Q52R^ channels were purified as previously described^58^, except after the total membrane fraction was prepared, membranes were re-suspended in buffer containing 0.2 M NaCl, 0.1 M KCl, 0.05 M HEPES pH 7.5, 4% trehalose, and 1 mg/mL brain PIP_2_ (Avanti Polar Lipids), and incubated at 4°C for 30 minutes before increasing the volume 10x in the same buffer without added lipids but including the detergent digitonin at a final concentration of 0.5%, then membranes were solubilized at 4°C for 90 minutes. As previously described^58^, after clarification by ultracentrifugation, the soluble fraction was incubated with anti-FLAG M2 affinity agarose for 10 hours and eluted at 4°C for 60 minutes with a buffer containing 0.2 M Na Cl, 0.1 M KCl, 0.05 M HEPES pH 7.5, 0.05% digitonin and 0.25 mg/mL FLAG peptide. Purified channels were eluted at ∼170 nM (∼150 μg/ml) and used immediately for cryo-EM grid preparation. K_ATP_ channel particles were also fixed and stained using Uranyl Acetate and sample quality was assessed by negative-stain-EM.

### CryoEM sample preparation and data acquisition

To increase protein adsorption to the cryoEM grids, and also to mitigate selective-orientation of K_ATP_ channel particles that occurs on commercial carbon surfaces, graphene-oxide ( GO) grids were prepared as previously described^59^. Briefly, gold Quantifoil R1.2/1.3 400 mesh grids were cleaned with acetone and glow-discharged for 60 seconds at 15 mA with a Pelco EasyGlow®, and 4 µL of 1 mg/mL Polyethylenimine (PEI, 40,000 MW) in 25 mM HEPES pH 7.9 was applied to each grid and incubated for 2 minutes followed by washing with water. Then, 0.1 mg/ml GO was vortexed vigorously and applied to the grid and incubated for 2 minutes followed by two washes with water. The GO grids were allowed to dry for 15 minutes and used within 2 hours for sample vitrification.

To prepare cryoEM samples, 3 µL of purified K_ATP_ channel complex was loaded onto fresh GO-coated grids for 30 s at 6 °C with a humidity of 100%. Grids were blotted for 2.5 seconds with a blotting force of −10 and cryo-plunged into liquid ethane cooled by liquid nitrogen using a Vitrobot Mark III (FEI).

Single-particle cryo-EM data was collected on a Titan Krios 300 kV cryo-electron microscope (ThermoFisher Scientific) in the Pacific Northwest CryoEM Center (PNCC), with a multi-shot strategy using beam shift to collect 27 movies per stage shift, assisted by the automated acquisition program SerialEM. Images were recorded on the Gatan K3 Summit direct-electron detector in super-resolution mode, post-GIF (20eV window), at 105,000x magnification (calibrated image pixel-size of 0.826 Å; super-resolution pixel size 0.413 Å); nominal defocus was varied between −1.0 and −2.5 µm across the dataset. The dose rate was kept around 24 e^-^/Å^2^/sec, with a frame rate of 35 frames/sec, and 78 frames in each movie ( i.e. 2.2 sec exposure time/movie), which gave a total dose of approximately 55 e^-^/Å^2^. Three grids that were prepared in the same session using the same protein preparation were used for data collection, and from these three grids, 2727, 3956 and 1576 movies (8,259 movies total) were recorded.

### CryoEM image processing

Super-resolution dose-fractionated movies were gain-normalized by inverting the gain reference in Y and rotating upside down, corrected for beam induced motion, aligned, and dose-compensated using Patch-Motion Correction in cryoSPARC2^60^ without binning. Parameters for the contrast transfer function (CTF) were estimated from the aligned frame sums using Patch-CTF Estimation in cryoSPARC2^60^ and binned by 2 with Fourier cropping. Micrographs were manually curated using sorting by curate exposures. The resulting 5241 dose-weighted motion-corrected summed micrographs were used for subsequent cryo-EM image processing. Particles were picked automatically using template-based picking in cryoSPARC2 based on 2D classes obtained from cryoSPARC live during data collection. For each of the three sets of data, particles were cleaned by three rounds of 2D classification in cryoSPARC2 ^60^. The combined particle stack contained 70,638 particles, which were then used for ab initio reconstruction in C1 requesting two classes. Only classes that had good alignments and contained full channel particles were included in subsequent rounds of classification. 40,896 particles from the best class were then subjected to ab initio reconstruction in C1 requesting three classes, which gave a class with only side, a class with only top/bottom, and a good class with 23,378 particles. These 23,378 particles were used for a final ab initio reconstruction using three classes, and gave 21,663 particles that sorted into a class with a straight transmembrane region (14,992 particles) and a class where the transmembrane region was more bent with the Kir6.2 core puckered up towards the extracellular space (6,657 particles). Further rounds of ab initio reconstruction resulted in equivalent classes without further improvement in particle classification. The particles were re-extracted using a 600 pixel box at 0.8265 Å/pix, duplicate particles within 20 Å from each other on the micrograph were removed, then used as input for non-uniform refinement^61^ in C1 with the 6,657 particle map reconstruction at 6.4 Å resolution and the 14,115 particle map reconstruction at 3.4 Å resolution. A non-uniform refinement in C4 symmetry imposed with the 14,115 particles resulted in a 2.9 Å resolution reconstructed cryoEM map using cryoSPARC auto mask tightening (Fig. S1). The auto-tightened FSC mask excluded the disordered NBD2 (Fig. S1c). Masks for FSC calculation that included the full K_ATP_ channel with or without micelle were generated using molmap in ChimeraX ^62^ (Fig. S1b,c), and resampled on the full map grid (600^3^ pixels). The FSC calculated using these static masks yielded 3.3 Å resolution for the full channel without micelle (Fig. S1b) and 3.5 Å resolution for the entire particle including the micelle (“Loose” in the FSC plot shown in Fig. S1b).

To assess conformational heterogeneity for the entire dataset, duplicate and corrupt particles were removed from the set of 70,638 particles from 2D classification to give 66,049 particles and refined using cryoSPARC homogeneous refinement with C4 symmetry imposed (4.0 Å resolution), then 4-fold symmetry expanded to give 264,196 particles (Fig. S2a). A mask covering the Kir6.2 tetramer, four TMD0 densities in SUR1, plus a large volume surrounding one of the SUR1 subunits, to have a large region covering all possible SUR1 locations, was generated using Chimera and used as a focused map to conduct 3D classification of symmetry expanded particles without particle alignment in cryoSPARC2. This yielded two dominant classes: class 1 ( 106,781 particles), which resembles the open structure derived using the workflow shown in Fig. S1a, and class 2 (53,201 particles), which has the Kir6.2^Q52R^ in the CTD-down position resembling previously published apo closed WT channel^38^. Particles from each of these classes were further classified into four classes using cryoSPARC 3D classification, and the classes with the most divergent conformations were compared (Fig. S2b-e). The reconstructed map resembling the published apo closed WT structure (PDB ID 7UQR) was well-fit by a rigid-body of that structure, but the resolution of the map was not sufficient to build a detailed atomic model.

To assess conformational heterogeneity within the high-resolution class of 14,115 particles of full-channel complexes that may have individual SUR1 subunits adopting independent conformations within a single K_ATP_ channel particle, particles were subjected to C4 symmetry expansion and 3D classification without particle alignment in cryoSPARC2 using the same mask described above. An initial focused 3D classification searching for 5 classes sorted particles into three dominant classes (class 0: 925 particles; class 1: 22474 particles; class 2: 10711 particles; class 3: 25664 particles; class 4: 194 particles). Class 1 to 3 showed three distinct SUR1 conformations, and subsequent local refinement using a loose mask covering the Kir6.2 tetramer, four TMD0 domains of SUR1, and one SUR1, resulted in map reconstructions of 3.12 Å, 3.99 Å, and 3.07 Å resolutions for class 1, 2, and 3, respectively (Fig. S6a,b). In all three of these particle classes, the NBDs are separated, but with substantial differences between the relative positions of NBD2, and all have better resolved maps corresponding to the NBD2 than for the overall consensus C4 non-uniform refinement.

In the C4 reconstruction of the full K_ATP_ channel particle, there is good density for nearly every side chain of Kir6.2 except for the N-terminal 31 residues and the C-terminal residues beyond residue 352. TMD0 and the transmembrane helicies of SUR1 also have clear density, with unclear density for the dynamic NBDs and loop regions, especially NBD2 which is poorly resolved due to apparent high flexibility when in the open conformation without NBD dimerization conditions.

To create an initial structural model, PDB ID 6BAA was fit into the reconstructed density for the full K_ATP_ channel particle using Chimera^63^, and then refined in Phenix^64^ as separate rigid bodies corresponding to TMD (32-171) and CTD 172-352) of Kir6.2 and TMD0/L0 (1-284), TMD1 (285-614), NBD1 (615-928), NBD1-TMD2-linker (992-999), TMD2 (1000-1319) and NBD2 (1320-1582). All sidechains and missing loops that had clear density were then built manually using Coot^65^. The resulting model was further refined using *Coot* and *Phenix* iteratively until the statistics and fitting were satisfactory (Table S1). Two models are presented. The first (PDB ID 8TI2) contains residues 32-352 for Kir6.2, and residues 1-1578 for SUR1 except for an extracellular loop (1043-1060), two loop regions (623-673 and 743-766) in NBD1 and the linker between NBD1 and TMD2 (928-986). The second model (PDB ID 8TI1) has the highly flexible NBD2 removed, and the SUR1 model ends at residue 1317. NBD1/2 and loop regions showed signs of disorder, thus many sidechains of residues in these regions were stubbed at Cβ. The N-terminal of Kir6.2, which in the closed NBD-separated conformation rests between TMD1 and TMD2 of SUR1, is not observed in this conformation.

In addition to protein density, N-acetylglucosamine (NAG), which is the common core of N-linked glycosylation, is modeled in the distinctively large density at the side chain of N10 in each SUR1 (Fig. 1b). Densities corresponding to two PIP_2_ molecules were also observed at the interface between Kir6.2 and SUR1 and were modeled after considering fits of other lipid molecules. Three other lipid sites per SUR1 monomer provided a useful contrast to the bulky density of the PIP_2_ head group (Fig. S3). The resulting model was further refined using *Coot*^65^ and *Phenix*^64,66,67^ iteratively until the statistics and fitting were satisfactory (Table S1). All structure figures were produced with UCSF Chimera^63^, ChimeraX^62^, and PyMol (http://www.pymol.org). Pore radius calculations were performed with HOLE implemented in *Coot* ^65^.

### Electrophysiology

For electrophysiology experiments, COSm6 cells were co-transfected with various combination of WT or mutant SUR1 and Kir6.2 cDNAs along with the cDNA for the Green Fluorescent Protein (to facilitate identification of transfected cells) using FuGENE® 6. Cells were plated onto glass coverslips twenty-four hours after transfection and recordings made in the following two days. All experiments were performed at room temperature as previously described ^68^. Micropipettes were pulled from non-heparinized Kimble glass (Fisher Scientific) on a horizontal puller (Sutter Instrument, Co., Novato, CA, USA). Electrode resistance was typically 1 −2 MΩ when filled with K-INT solution containing 140 mM KCl, 10 mM K-HEPES, 1 mM K-EGTA, 1 mM EDTA, pH 7.3. ATP was added as the potassium salt. Inside-out patches of cells bathed in K-INT were voltage-clamped with an Axopatch 1D amplifier (Axon Inc., Foster City, CA). ATP or porcine brain PIP_2_ (Avanti Polar Lipids; prepared in K-INT and bath sonicated in ice water for 30 min before use) were added to K-INT as specified in the figure legend. All currents were measured at a membrane potential of −50 mV (pipette voltage = +50 mV). Initial currents prior to PIP_2_ exposure shown in Fig. 3 were calculated by subtracting currents measured in K-INT with 1 mM ATP (which inhibits >99% of K_ATP_ currents) from currents in K-INT solution to account for leak currents. Data were analyzed using pCLAMP10 software (Axon Instrument). Off-line analysis was performed using Clampfit and GraphPad. Data were presented as mean ± standard error of the mean (S.E.M). ATP inhibition dose-response curves were obtained by fitting data to the Hill equation (I_rel_ = 1/(1 + ([ATP]/IC_50_)^H^)), where I_rel_ is the current relative to the maximum currents in K-INT solution (expressed as % current in Figs. 7c, S7a), IC_50_ is the ATP concentration that causes half-maximal inhibition, and H is the Hill coefficient. Note H was allowed to vary for the curves shown in Figs. 7c, S7a.

### Rb^+^ efflux assay

COSm6 cells were transiently transfected with various combination of WT or mutant SUR1 and Kir6.2 cDNAs using FuGENE® 6. Untransfected cells were included as background control. Cells were cultured in medium containing 5 mM RbCl overnight. The next day, cells were washed quickly twice in Ringer’s solution (5.4 mM KCl, 150 mM NaCl, 1 mM MgCl_2_, 0.8 mM NaH_2_PO_4_, 2 mM CaCl_2_, 25 mM HEPES, pH 7.2) with no RbCl. Rb efflux was measured by incubating cells in Ringer’s solution (for experiments shown in Fig. 7d) or Ringer’s solution supplemented with 5 mM glucose (for experiments shown in Fig. S7b) over a 40 min period. Inclusion of 5 mM glucose increases intracellular ATP/ADP ratios to suppress channel activity, allowing for better detection of channels showing mild gain-of-function phenotype as shown in some of the mutant channels in Fig. S7b). At the end of the 40 min incubation, Efflux solution was collected and cells lysed in Ringer’s solution plus 1% Triton X-100. Rb concentrations in both the efflux solution and cell lysate were measured using an Atomic Adsorption Instrument Ion Channel Reader (ICR) 8100 from Aurora Biomed. Percent efflux was calculated by dividing Rb in the efflux solution over total Rb in the efflux solution and cell lysate. For each experiment, duplicates were included as technical repeats and the average taken as the experimental value. For all channel combinations, 3-6 separate transfections were carried out in parallel as biological repeats. Data were presented as mean ± standard error of the mean (S.E.M) and statistical analysis performed by one-way ANOVA with Tukey’s post hoc test in GraphPad.

## Data availability

The cryo-EM maps have been deposited in the Electron Microscopy Data Bank (EMDB), and the coordinates have been deposited in the PDB under the following accession numbers: PDB ID 8TI2 and EMD-41278 (NBD2 modeled as main chain atoms only); PDB ID 8TI1 and EMD-41277 (NBD2 not modeled).

## Author contributions

CMD designed and performed experiments, analyzed data, prepared figures, wrote and edited the manuscript. YYK performed electrophysiology and Rb efflux experiments, analyzed data, prepared figures, and edited the manuscript. PZ analyzed data, prepared figures and edited the manuscript. AE performed Rb efflux experiments, prepared figures and edited the manuscript. SLS conceived the project, performed electrophysiology experiments, prepared figures, wrote and edited the manuscript.

## Acknowledgements

A portion of this research was supported by NIH grant U24GM129547 and performed at the Pacific Northwest Cryo-EM Center (PNCC) at Oregon Health and Science University and accessed through EMSL (grid.436923.9), a DOE Office of Science User Facility sponsored by the Office of Biological and Environmental Research, with special thanks to Dr. Nancy Meyer for help with cryoEM data collection. We acknowledge the support by the National Institutes of Health grant R01DK066485 (to SLS) and the OHSU Tartar trust foundation ( to CMD). We are grateful to Zhongying Yang for help preparing adenovirus constructs and John Allen for preparing plasmids. We thank Drs. Min Woo Sung, Katarzyna Ciazynska, and Bruce L. Patton for helpful discussions and Dr. Bruce L. Patton for comments on the manuscript.

---

**Figure S1.**
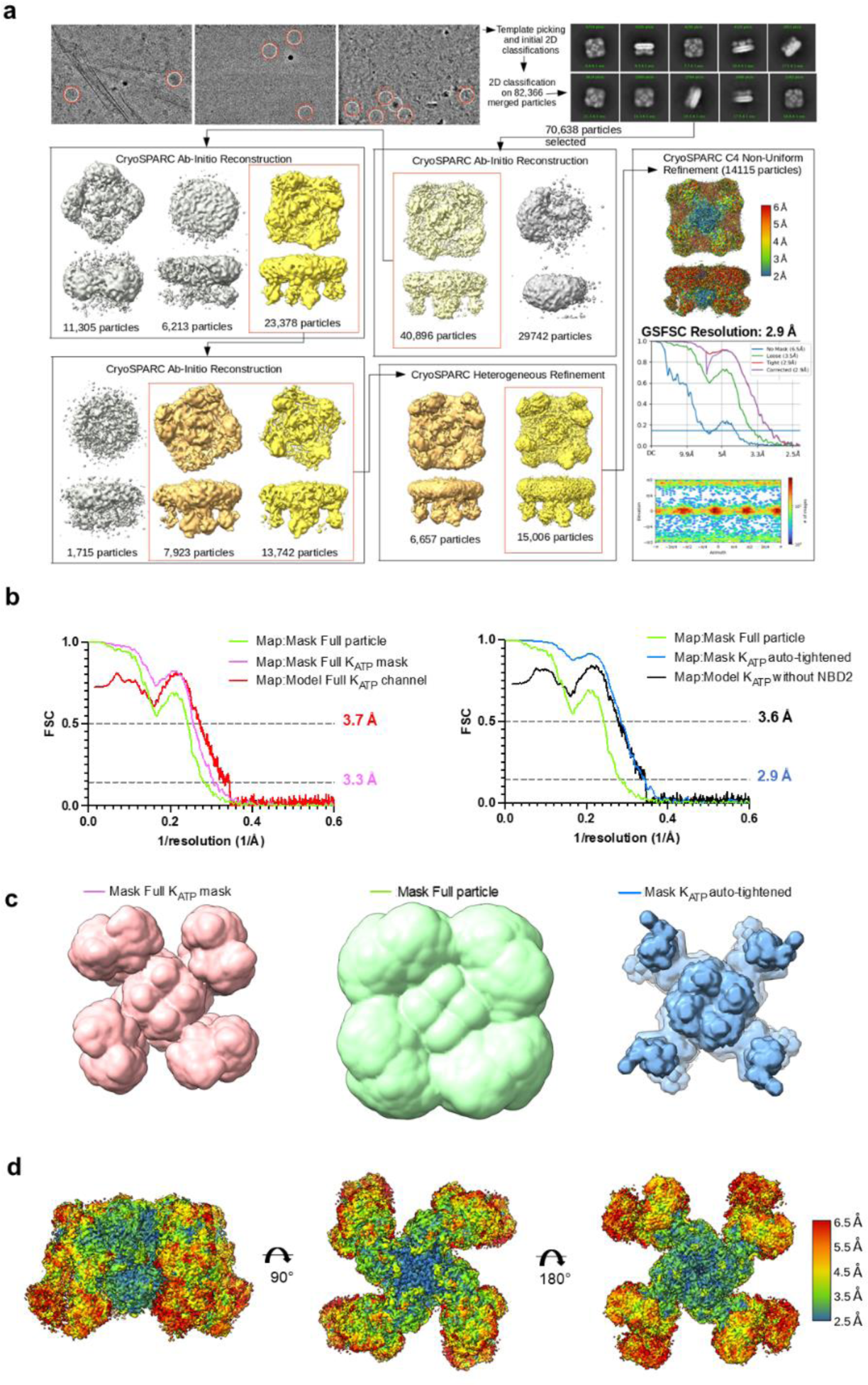
CryoEM data processing. **(a)** Workflow showing the total number of particles yielding the 2.9 Å C4 Non-uniform refinement reconstruction is 14,115 (FSC auto mask for GSFSC calculation). The three raw images on the top left corner serve to illustrate the particles on GO layer (GO edges are seen in the left micrograph) selected (red circles) for processing **(b)** Fourier Shell Correlation (FSC) curves between two independent half-maps calculated within masks shown in (c), and for model:map (unmasked) for the full K_ATP_ channel model and map (PDB ID 8TI2; EMD-41278) (*left*), and for the K_ATP_ channel without NBD2 (PDB ID 8TI1) (*right*). **(c)** Masks used for FSC calculations: full particle including micelle (green), a mask for full K_ATP_ channel without the micelle (pink), and a mask created by cryoSPARC auto-mask tightening that excludes NBD2 (blue), shown in cytoplasmic view. **(d)** The local resolution estimate for the reconstructed map at 4 σ (0.08 V) contour, with micelle density not shown, in side view (*left*), top view (*middle*) and bottom view (*right*). No local filtering or local sharpening was used for visualization. Local resolution estimates were calculated in cryoSPARC and visualized in ChimeraX. FSC curves were calculated in PHENIX and cryoSPARC.

**Figure S2.**
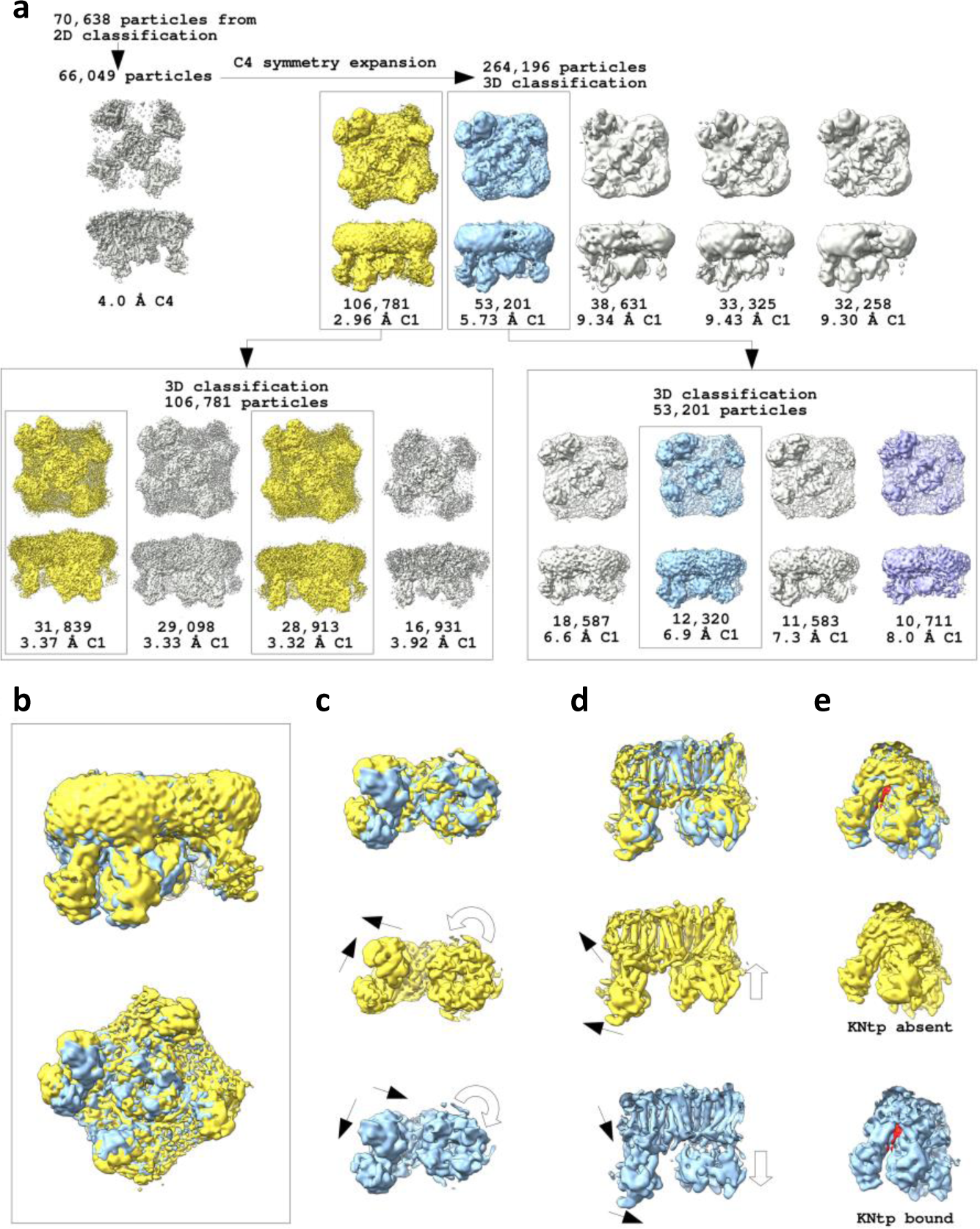
Distinct conformation classes revealed by symmetry expansion and focused 3D classification. **(a)** Workflow to obtain distinct conformation classes (details described in Methods). **(b)** Overlay of full channel map generated by combining the two yellow classes from the second round of 3D classification (shown in yellow) and the blue map from the second round of 3D classification(shown in blue) viewed from the side and the bottom. The yellow map resembles the PIP_2_-bound open structure derived from C4 non-uniform refinement (Figs. 1a, S1a), while the blue map resembles previously published closed apo WT structure (PDB ID 7UQR), and will be referred to as apo SUR1/Kir6.2^Q52R^ structure. **(c)** Cytoplasmic view of the Kir6.2^Q52R^ tetramer plus one SUR1 in the two maps superimposed (top) or separately. In the blue map, Kir6.2^Q52R^-CTD is rotated clockwise relative to the yellow map. Also in the blue map, the NBDs of SUR1 are closer to the Kir6.2^Q52R^ tetramer and SUR1 is rotated opposite of Kir6.2^Q52R^-CTD, compare to the yellow map. **(d)** Side view of the maps in (c) showing Kir6.2^Q52R^-CTD is docked up to the membrane in the yellow map, and extended down from the plasma membrane in the blue map. Also, SUR1 is tilted away from the Kir6.2^Q52R^ tetramer in the yellow map, but tilted towards the Kir6.2^Q52R^ tetramer in the blue map. The curved and straight open arrows mark the relative rotation and translation from the membrane of the Kir6.2^Q52R^-CTD. The solid arrows indicate relative movements of the SUR1. **(e)** Side view of the SUR1 ABC core showing the presence of cryoEM density corresponding to the Kir6.2^Q52R^ N-terminal peptide (KNtp, red density) in the apo closed SUR1/Kir6.2^Q52R^ map (blue) but absence of KNtp density in open SUR1/Kir6.2^Q52R^ map (yellow).

**Figure S3.**
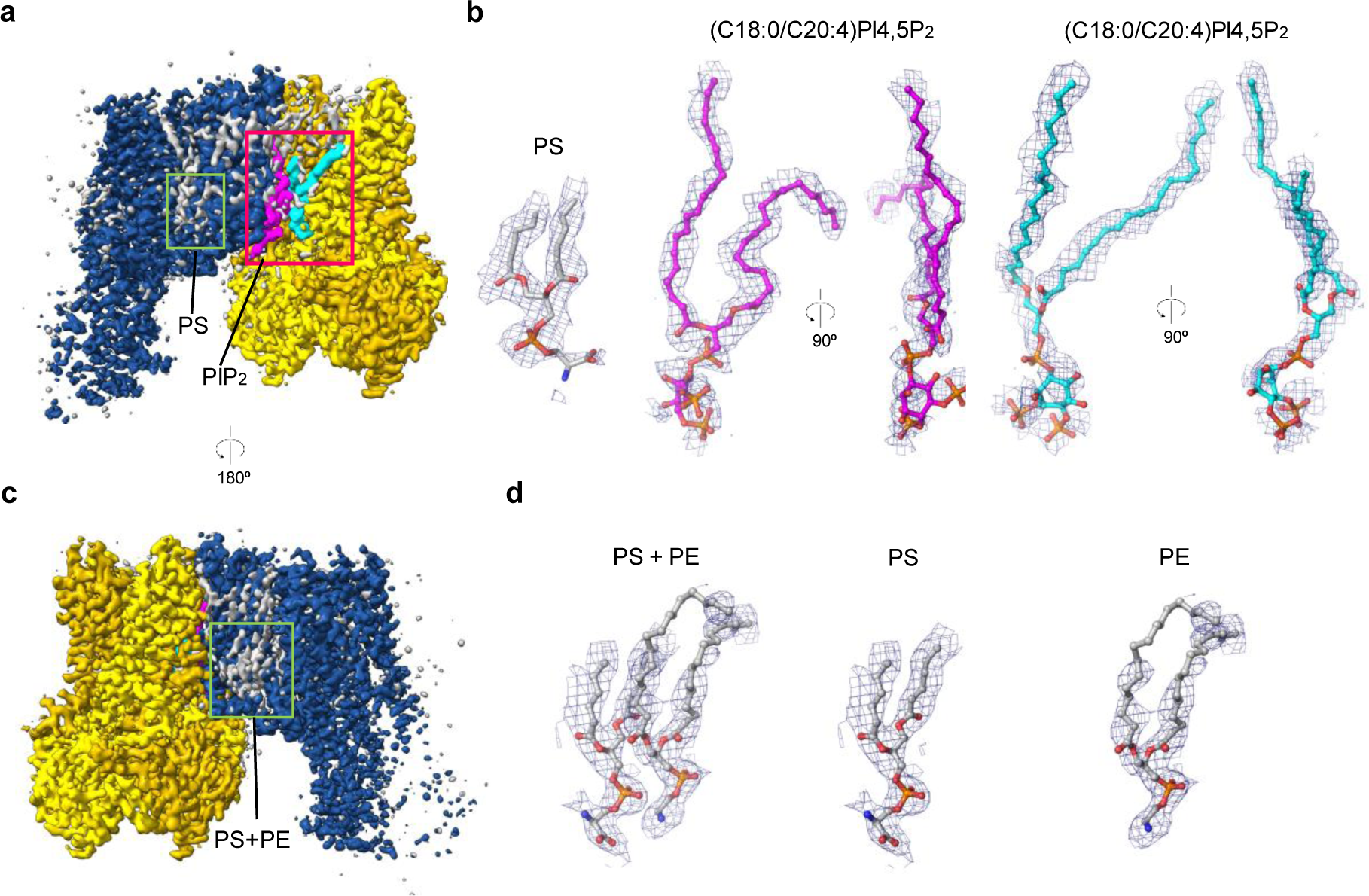
CryoEM lipid density fitting. **(a)** CryoEM map reconstructed from symmetry expansion of the ∼14k particles followed by local refinement with a focused mask of Kir6.2 tetramer plus one SUR1, with density modification in Phenix and map segmentation in Chimera, shows the cryoEM densities of SUR1 and the Kir6.2 tetramer colored blue and yellow, respectively. Two PIP_2_ molecules are shown in magenta and cyan, and other density including lipid density shown in grey (0.08V/4σ contour). **(b)** Isolated lipid density located in the inner leaflet between SUR1-TMD0 and SUR1-TMD1 fit with a model of phosphatidylserine (PS, grey carbons, 0.08V/4σ contour) and isolated lipid density at each of the two PIP_2_ binding sites located between SUR1-TMD0 and Kir6.2 shown fit with model of the first-(cyan carbons) and second-(magenta carbons) (C18:0/C20:4) PI(4,5)P_2_ in two different rotational views. **(c)** Rotation of the map 180° relative to panel A reveals other densities (grey, 0.08V/4.0σ contour) corresponding to two adjacent lipids primarily associated with the TMD0 domain. **(d)** Adjacent lipid densities can be well fit as a phosphatidylserine (PS) and phosphatidylethanolamine (PE), with the amine group of PE hydrogen bonding to the phosphate group of PS. On the right, density for each of these adjacent lipid molecules is isolated to show the fit of the lipid model into the lipid’s corresponding density. These five lipid densities (20 lipids for full-channel) that are captured in the micelle are closely associated withthe K_ATP_ channel and are sufficiently resolved to allow modeling. Note lipid or detergent densities are also observed in the outer leaflet space; however, they are not sufficiently resolved to allow modeling.

**Figure S4.**
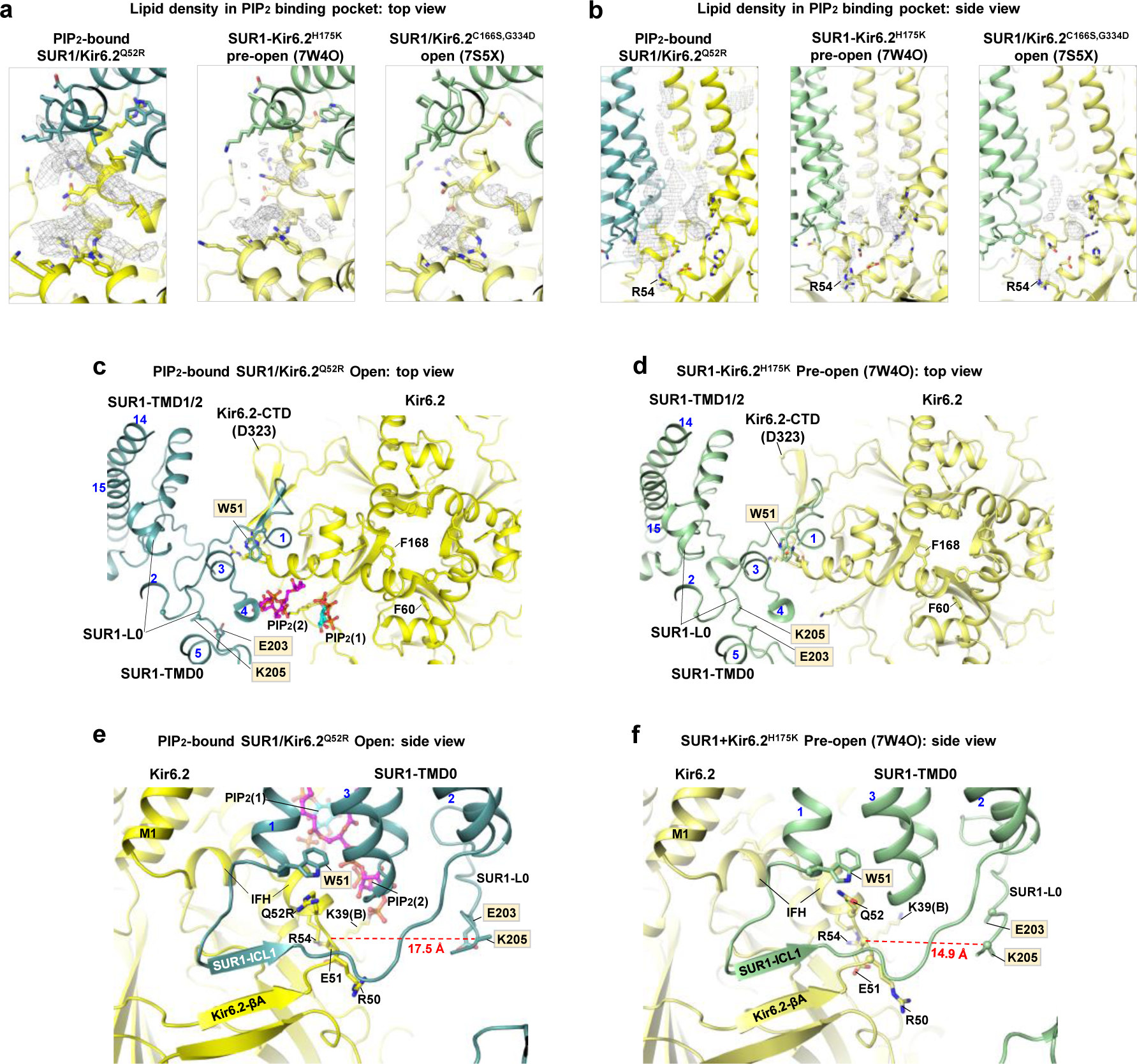
Comparison of PIP_2_-bound open K_ATP_ channel structure with other open K_ATP_ structures. **(a, b)** Comparison of the lipid density (grey mesh) in the PIP _2_ binding pocket in the PIP_2_-bound SUR1/Kir6.2^Q52R^ structure, SUR1-Kir6.2^H175K^ fusion channel pre-open structure (in the presence of diC8-PIP_2_; EMD-32310, PDB ID 7W4O), and SUR1/Kir6.2^C166S,G334D^ open structure (no PIP_2_ added; EMD-24842, PDB ID 7S5X), viewed from the top (a) and the side (b). In (b), the cryoEM density for Kir6.2 R54 is shown as a reference. **(c, d)** Top view of PIP_2_-bound SUR1/Kir6.2^Q52R^ open channel (c) and SUR1-Kir6.2^H175K^ fusion pre-open channel (PDB ID 7W4O) (d), showing similarities in side chain orientations for Kir6.2 F60 and F168 (the gate residue at the helix bundle crossing) and SUR1-W51, and rotation position of the Kir6.2 cytoplasmic domain (CTD) marked by residue D323. **(e, f)** Comparison of the two structures in (c) and (d) viewed on the side, showing close proximity of SUR1-W51 to Kir6.2-Q52R (e) or Q52 (f), as well as the widening of the ATP binding pocket marked by the distance (dashed red line) between the Cα atoms of Kir6.2-E51 and SUR1-K205.

**Figure S5.**
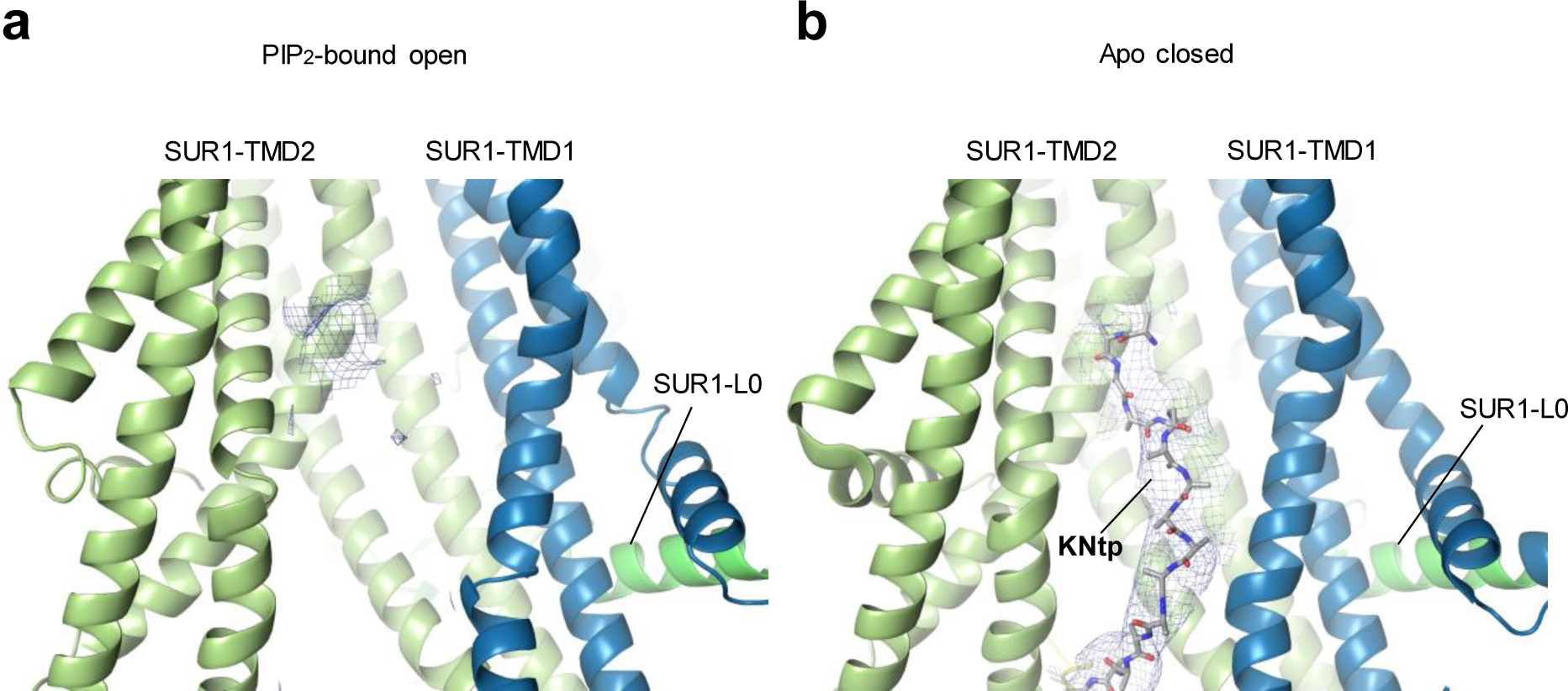
Comparison of KNtp density in the SUR1-ABC core in the PIP_2_-bound open versus apo closed SUR1/Kir6.2^Q52R^ structures. **(a)** In the PIP_2_-bound SUR1/Kir6.2^Q52R^ open channel, no KNtp density is observed, consistent with the absence of the Kir6.2^Q52R^-Ntp or a highly flexible Kir6.2^Q52R^-Ntp. Note the density protruding from SUR1-TMD2 TM helix corresponds to W1297 with some R1145 contribution. **(b)** In the apo SUR1/Kir6.2^Q52R^ closed channel, a clear peptide density corresponding to Kir6.2^Q52R^-Ntp is clearly present. For density shown in the KNtp binding cleft in SUR1, both maps were filtered to 7 Å and contoured to 6.5 σ.

**Figure S6.**
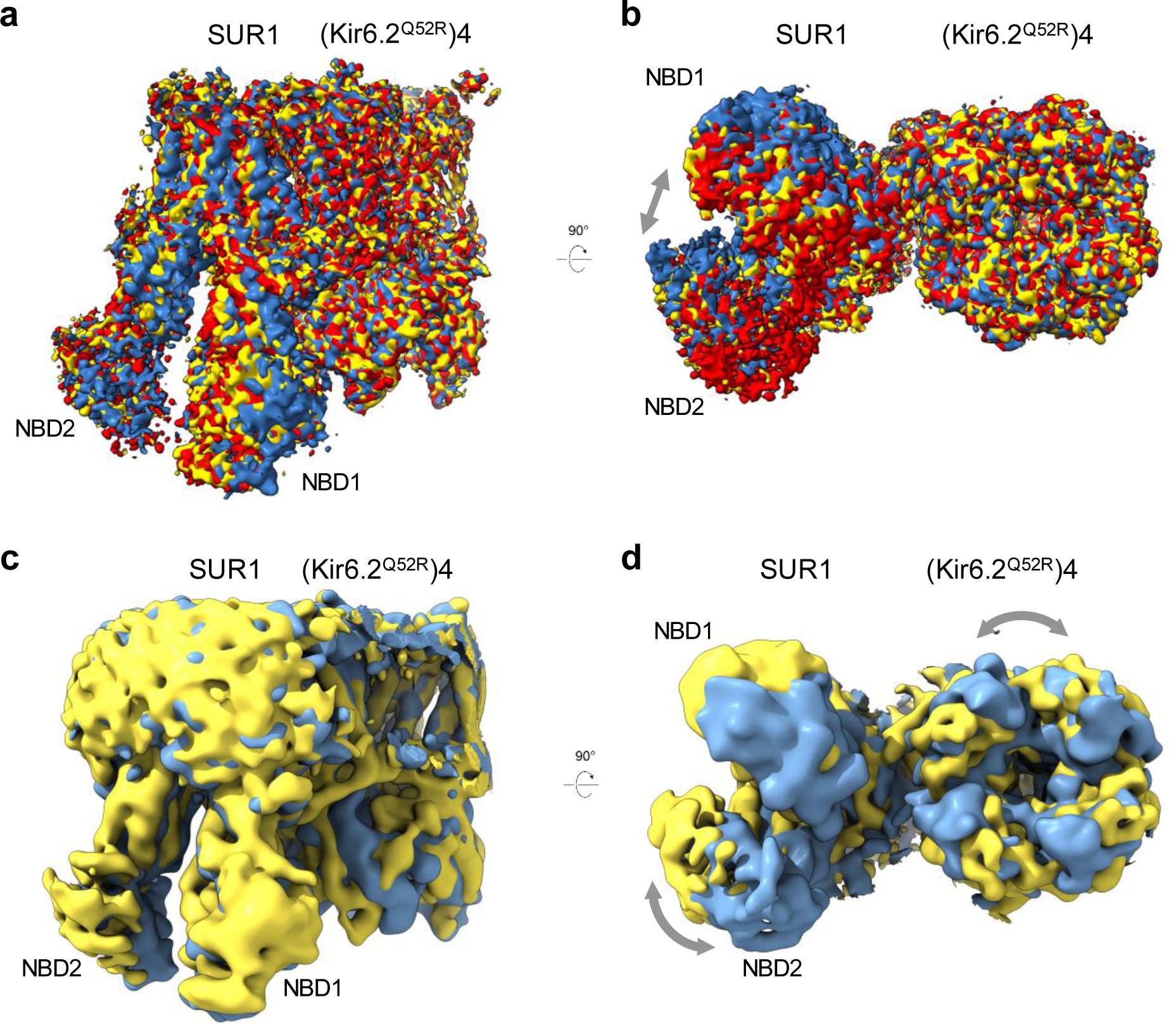
Conformational dynamics of the SUR1 subunit revealed by CryoSPARC 3D classification. **(a,b)** Dynamic range of SUR1 within the final open particle class (Fig. S1) was determined by symmetry expansion followed by 3D classification. The 14,115 particles within the final open class gave 56,460 particles, which were subjected to 3D classification with a focused mask of Kir6.2 tetramer plus one SUR1. Three unique NBD-separated SUR1 positions were resolved into three classes. Reconstructed maps of class 1 (3.0 Å resolution, 23,454 particles, red map), class 2 (3.3 Å resolution, 16,863 particles, yellow map) and class 3 (3.4 Å resolution, 13,604 particles, blue map) reconstructed from a local non-uniform refinement with density modification in Phenix. (a) Transmembrane view shows the variability in SUR1 including the transmembrane regions of TMD1 and TMD2, in an open Kir6.2 tetramer position. (b) View from the cytoplasmic side highlights the difference in the SUR1-NBD positions within the open particle class, with mobility seen in both SUR1-NBD1 and SUR1-NBD2. The grey arrow indicates the direction of the NBD position differences. **(c,d)** The reconstructed maps for the PIP_2_-bound open class (yellow, 7.0 Å filtered resolution) and apo closed class (blue, 7.0 Å filtered resolution) within the SUR1/Kir6.2^Q52R^ dataset (workflow in Fig. S2), show the relative SUR1 and Kir6.2 domain positions as the channel transitions between open and closed conformations. (c) Transmembrane view shows the difference in the SUR1-NBD position and the rotation of the Kir6.2-CTD. (d) View from the cytoplasmic side highlights the difference in the SUR1-NBD positions relative to the Kir6.2 subunit, with the open conformation having the SUR1-NBD further away from the Kir6.2 cytoplasmic domain which is rotated counterclockwise relative to the closed position. The grey curved arrows indicate the rotational position difference in Kir6.2^Q52R^-CTD and SUR1-NBDs

**Figure S7.**
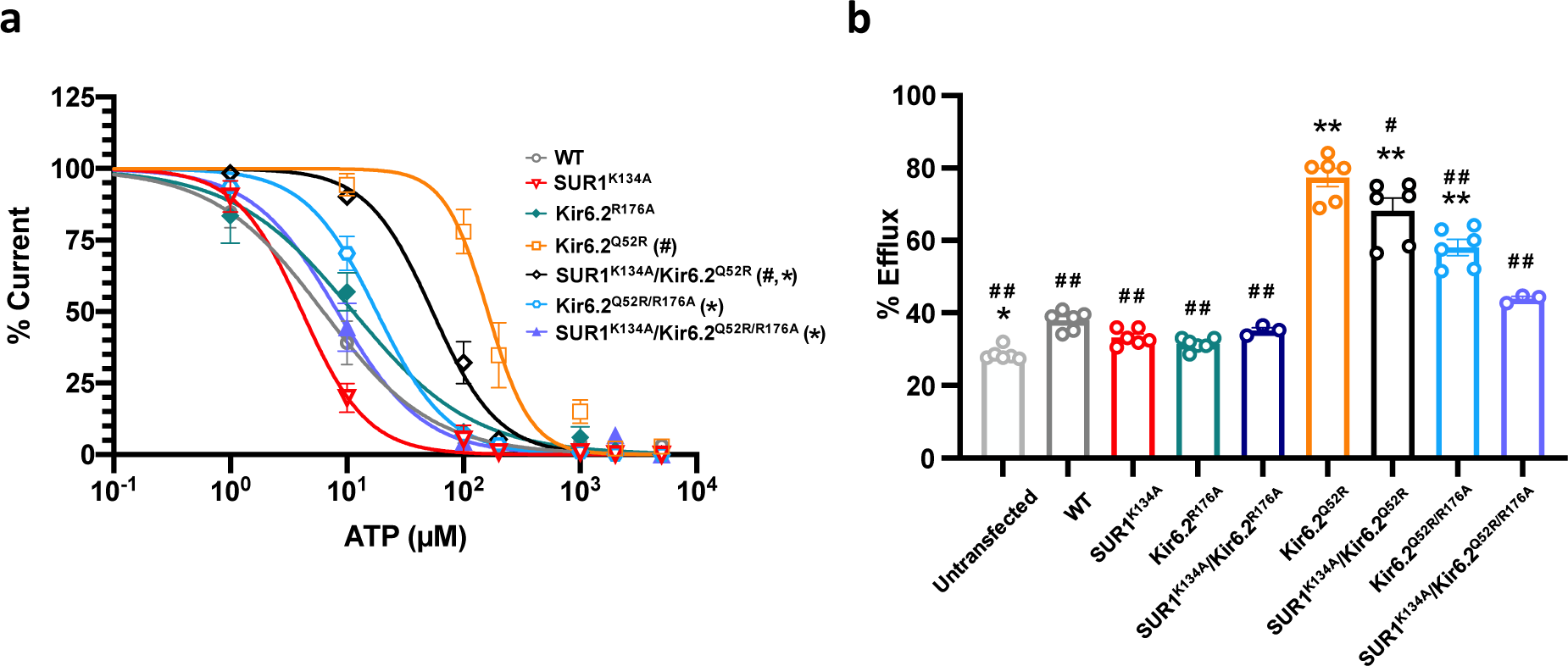
Perturbation of PIP_2_ binding residues diminished the gain-of-function effect of Kir6.2^Q52R^. **(a)** ATP dose response of WT (SUR1/Kir6.2) channels, and channels containing the PIP_2_ binding site mutations SUR1^K134A^ (SUR1^K134A^) or Kir6.2^R176A^, as well as the gain-of-function disease mutation Kir6.2^Q52R^alone or in combination with single PIP_2_ binding site mutations (SUR1^K134A^/Kir6.2^Q52R^, or Kir6.2^Q52R/R176A^) or double PIP_2_ binding site mutations (SUR1^K134A^/Kir6.2^Q52R/R176A^). Curves were obtained by fitting the data points (mean ± SEM of 3-6 patches) to the Hill equation (see Methods). ^#^*p* < 0.001. compared to WT channel; \**p* < 0.001, compared to SUR1/Kir6.2^Q52R^ channel (Kir6.2^Q52R^), by one-way ANOVA with Tukey’s *post hoc* test. **(b)** Rb^+^ efflux of various channels expressed in COSm6 cells measured in Ringer’s supplemented with 5 mM glucose. Untransfected cells were included to show background efflux. \**p* < 0.05, \*\**p* < 0.0001 compared to WT; ^#^*p* < 0.05, ^##^*p* < 0.0001 compared to Kir6.2^Q52R^, n=3-6; one-way ANOVA with Tukey *post hoc* test).

**Figure S8.**
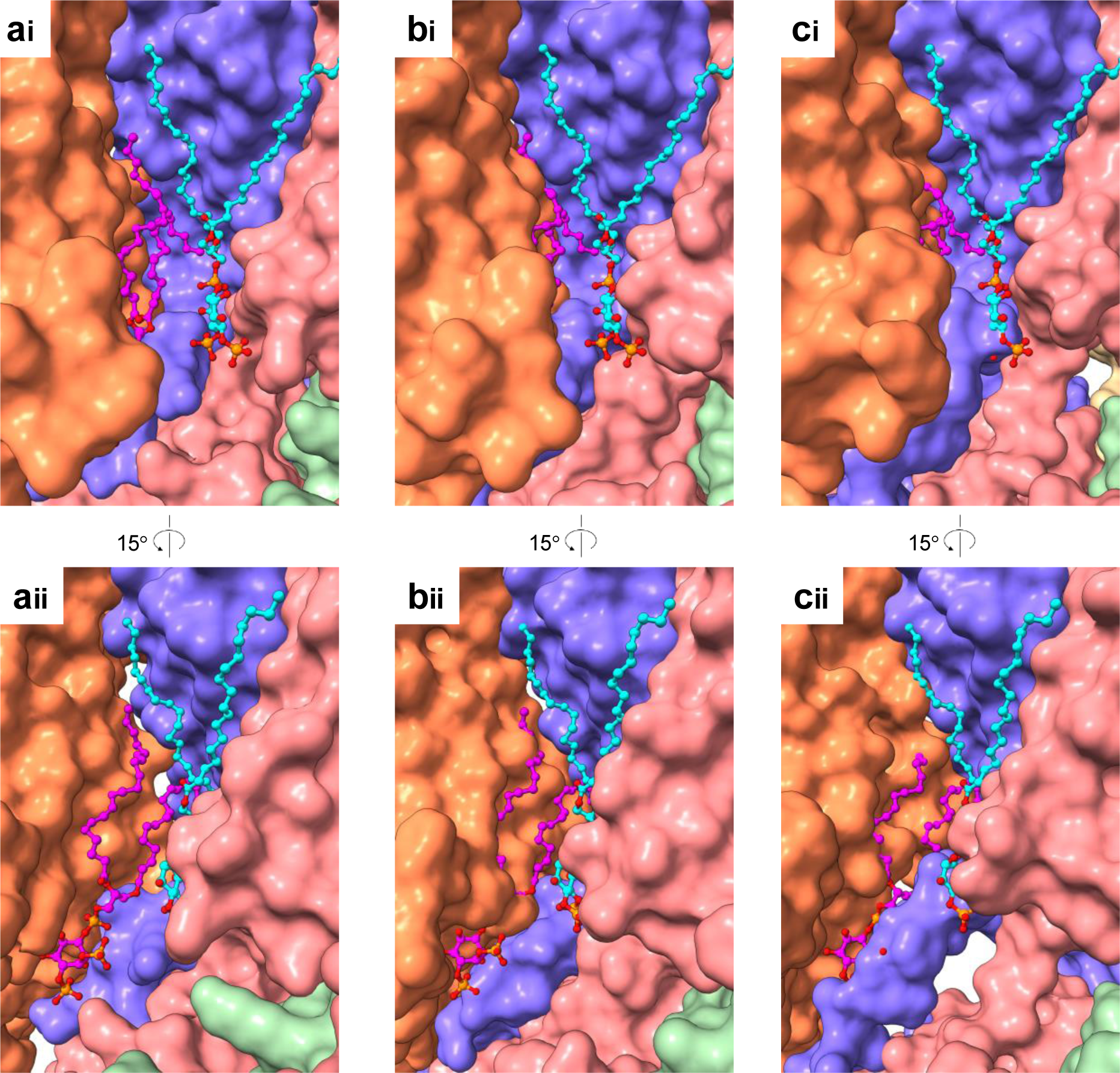
The K_ATP_ channel closed conformations clash with the novel second PIP_2_ molecule, but both the open and closed conformations could accommodate the conserved first PIP _2_. **(a)** The open SUR1/Kir6.2^Q52R^ K_ATP_ channel with the conserved first PIP_2_ (cyan carbons) and novel second PIP_2_ (magenta carbons) shown interacting with Kir6.2 subunits (pink, green and blue surface) and the SUR1 subunit (orange surface), with the two panels rotated ∼15° to show the surface of each PIP_2_ site. **(b)** The closed K_ATP_ channel bound to repaglinide and ATP (PDB ID 7TYS, with Kir6.2-CTD in the up position) aligned with the PIP_2_ molecules of the open channel shows that the closed K_ATP_ channel accommodates the conserved first PIP_2_ but clashes with the novel second PIP_2_. **(c)** The apo closed SUR1/Kir6.2^Q52R^ K_ATP_ channel (with Kir6.2-CTD in the down position, similar to previously reported apo WT structure PDB ID 7UQR; see Fig. S2) also shows clash with PIP_2_ at the second site but not the first site.

**Table S1.**
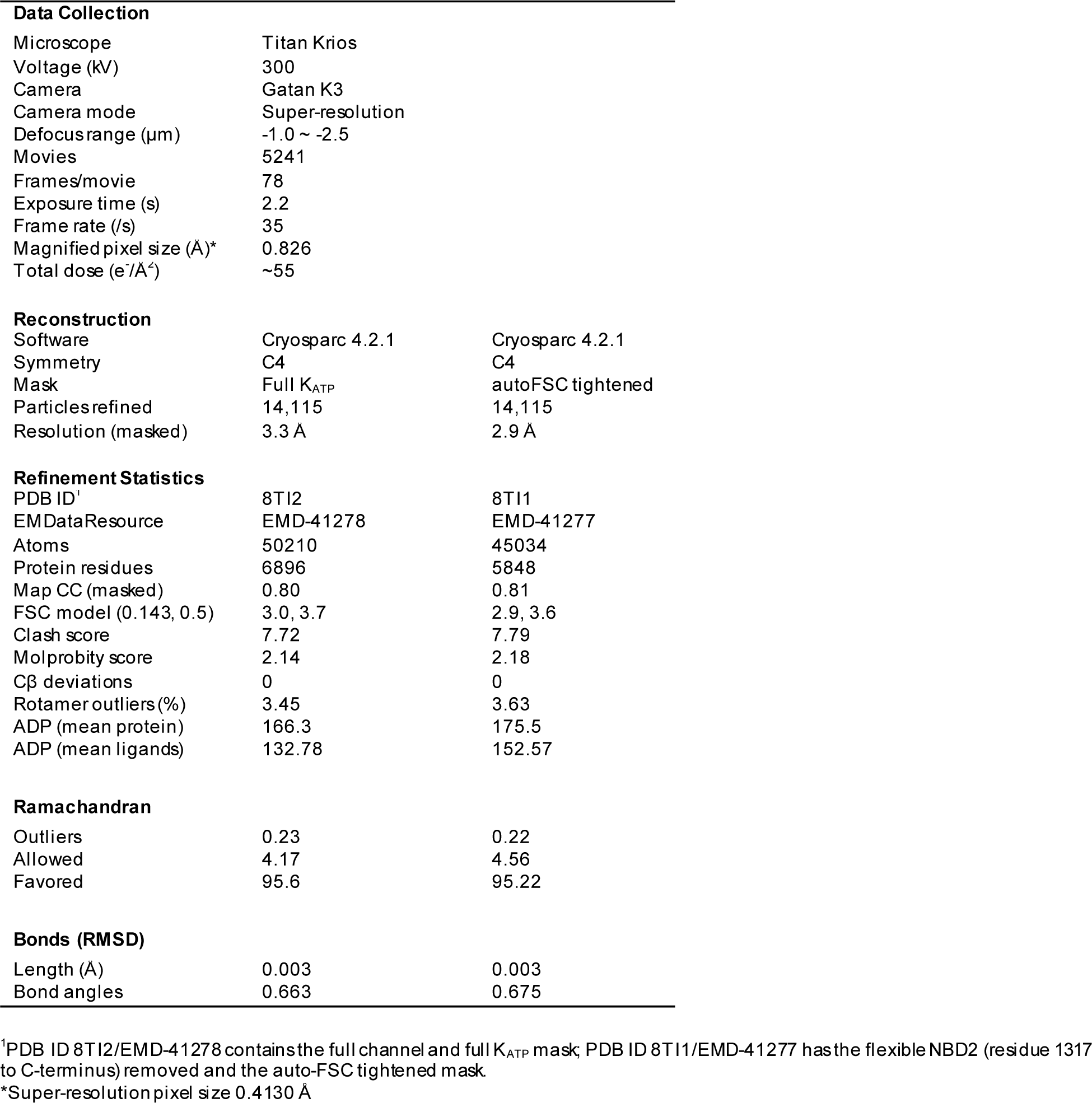
Model statistics of the PIP_2_-bound SUR1/Kir6.2^Q52R^ structure.

**Movie 1. Morph between PIP_2_-bound open SUR1/Kir6.2^Q52R^ channel structure and ATP/Repaglinide-bound closed WT channel structure.**

